# Universally available herbal teas based on sage and perilla elicit potent antiviral activity against SARS-CoV-2 variants of concern by HMOX-1 upregulation in human cells

**DOI:** 10.1101/2020.11.18.388710

**Authors:** Vu Thuy Khanh Le-Trilling, Denise Mennerich, Corinna Schuler, Roman Sakson, Julia K. Lill, Dominik Kopczynski, Stefan Loroch, Yulia Flores-Martinez, Benjamin Katschinski, Kerstin Wohlgemuth, Matthias Gunzer, Folker Meyer, Ulf Dittmer, Albert Sickmann, Mirko Trilling

## Abstract

The current SARS-CoV-2/COVID-19 pandemic wreaks medical and socioeconomic havoc. Despite the availability of vaccines, cost-effective acute treatment options preventing morbidity and mortality are urgently needed. To identify affordable, ubiquitously available, and effective treatments, we tested herbs consumed worldwide as herbal teas regarding their antiviral activity against SARS-CoV-2. Aqueous infusions prepared by boiling leaves of the *Lamiaceae* perilla and sage elicit potent and sustained antiviral activity against SARS-CoV-2 in therapeutic as well as prophylactic regimens. The herbal infusions exerted antiviral effects comparable to interferon-β and remdesivir but outperformed convalescent sera and interferon-α2 upon short-term treatment early after infection. Based on protein fractionation analyses, we identified caffeic acid, perilla aldehyde, and perillyl alcohol as antiviral compounds. Global mass spectrometry (MS) analyses performed comparatively in two different cell culture infection models revealed changes of the proteome upon treatment with herbal infusions and provided insights into the mode of action. As inferred by the MS data, induction of heme oxygenase 1 (HMOX-1) was confirmed as effector mechanism by the antiviral activity of the HMOX-1-inducing compounds sulforaphane and fraxetin. In conclusion, herbal teas based on perilla and sage exhibit antiviral activity against SARS-CoV-2 including variants of concern such as Alpha, Beta, Delta, and Omicron.

## Introduction

Fossil records suggest that humans may have applied plants as medicine at least from the Middle Palaeolithic age some 60,000 years ago ^1,2^. Across cultures and spanning thousands of years, humans consumed aqueous plant infusions as teas. The first textual reference to tea consumption dates back to 59 years before the current era (BCE) and physical evidence even dates back to 255±80 years BCE ^3^. In addition to reasons of enjoyment and taste, teas are frequently applied for disease prophylaxis, therapy, or symptom alleviation. A major distinction is made between genuine teas based on *Camellia sinensis* infusions versus various types of herbal teas. For the latter, parts of other plants are boiled in water generating complex aqueous infusions. Especially members of the *Lamiaceae* family comprising sage (*Salvia officinalis*) and perilla (*Perilla frutescens*) are ubiquitously used to prepare herbal teas. Additionally, important spices such as thyme, basil, mint, rosemary, marjoram, oregano, and lavender also belong to the *Lamiaceae*. Across the world, the edible plant perilla and its variations have a variety of names such as *Tía tô* (Vietnam), *rattlesnake weed* (US), *silam* (India and Nepal), *shiso* and *egoma* (Japan), *deulkkae* (Korea), *zĭsū* and *sūzĭ* (China) highlighting its broad distribution. In parts of Japan, it is referred to as *jūnen* (“ten years”) since it is believed to be able to add ten years to a person’s lifespan. Members of the *Lamiaceae* family are well described for their medicinal effects against various diseases including pneumonia and cough ^4^. While perilla is very popular in Asia, the related plant sage is more common in Europe and America. The name *Salvia officinalis* already indicates its medicinal potential since the *officinal* was a room in a monastery dedicated to herbs and medicine. Intriguingly, perilla and sage extracts indeed possess antimicrobial activities (e.g., ^5,6^) In the era of modern medicine, some people have reservations concerning the use of traditional and herbal medicines. However, a highly relevant fraction of recently approved modern therapeutics directly or indirectly originate from natural products ^7^ - some of which known to and applied by our ancestors. In this respect, the antimalarial lactone artemisinin derived from the sweet wormwood (*Artemisia annua*) is amongst the best-known examples ^8,9^.

Humans have been exposed to coronaviruses (CoV) for ages, given their broad prevalence in mammals (e.g., bats) and birds. At least seven CoVs are capable of autochthonous propagation in the human population: human CoV (HCoV)-HKU1, HCoV-NL63, HCoV-229E, HCoV-OC43, HCoV-SARS, HCoV-MERS, and the severe acute respiratory syndrome coronavirus 2 (SARS-CoV-2). The latter causes the current global pandemic of coronavirus disease 2019 (COVID-19). SARS-CoV-2 was first recognized and genetically defined in Wuhan, China ^10,11^. In various aspects, it shows similarities to SARS-CoV-1, however, it also exhibits certain specialties ^12,13^ such as the capacity of very efficient replication in the upper respiratory tract and the corresponding efficacy of human-to-human transmission. Given the broad coverage of this topic, we refer the reader to review articles concerning SARS-CoV-2 and COVID-19 (e.g., ^14-18^). At the time of writing, more than 475 million individuals experienced laboratory-confirmed SARS-CoV-2 infections and over 6 million people succumbed in the context of COVID-19. According to the Johns Hopkins dashboard ^19^, all relevant countries and regions are affected by SARS-CoV-2 cases - several of which are developing nations with very limited resources for the medical sector, especially when faced with overwhelming numbers of infected individuals ^20^. Since the onset of the COVID-19 pandemic in December 2019, SARS-CoV-2 has continuously evolved, with numerous variants emerging across the world. Depending on prevalence and clinical and epidemiological characteristics, these variants are classified as “variant of interest” (VOI), “variant under monitoring” (VUM), and “variant of concern” (VOC). As of March 2022, there are five SARS-CoV-2 lineages designated as VOCs (Alpha, Beta, Gamma, Delta, and Omicron variants). Delta was the most prevalent variant in 2021, but in 2022, Omicron has overtaken Delta as the predominant variant (https://nextstrain.org/ncov/gisaid/global). Certain VOCs exhibit increased transmissibility compared to the original virus and others the potential to increase disease severity. Moreover, some VOCs show decreased susceptibility to vaccine- and infection-induced immune responses, and thus possess an elevated ability to reinfect previously infected and recovered as well as vaccinated individuals.

A variety of SARS-CoV-like viruses can be found in bats, indicating an alarming reservoir of coronaviruses that could cause zoonotic animal-to-human spill over transmissions ^21-23^. Beyond the alpha- and beta-coronaviruses, which are known to comprise human coronaviruses, recent data indicate that humans can also acquire other coronaviruses such as porcine delta-coronaviruses from farm animals ^24^. Based on the facts that CoVs are present in animals such as rodents, bats, pigs, and cats residing in utmost proximity of human settlements and civilization and seem to have caused human epidemics in the past ^25,26^, we speculated that human culture might provide certain behavioral adaptations to coronavirus infections. Such knowledge may be applicable to alleviate some of the hardship and suffering caused by SARS-CoV-2 in a process of “cultural repurposing”. People with respiratory diseases often consume herbal products and teas in attempts of self-medication. Faced with the COVID-19 pandemic, people reported that they have changed their behavior accordingly. In two studies comprising thousands of people, up to 57.6% of individuals reported having used nutritional supplements or herbal products, usually as teas, in attempts to protect themselves from COVID-19 ^27,28^. Therefore, we wondered how effective herbal teas actually are against SARS-CoV-2 including VOCs.

## Results

### Perilla and sage contain water-soluble heat-stable components active against SARS-CoV-2 replication

Since efficient and affordable treatment options for COVID-19 patients are still not sufficiently available, we aimed to determine the therapeutic potential of herbal teas. To evaluate two universally available *Lamiaceae* commonly used in traditional medicine, perilla and sage, in terms of their ability to elicit antiviral activity against SARS-CoV-2, we applied an experimental setup that reflects short-term incubation of infected cells with herbal teas. We infected highly SARS-CoV-2-permissive Vero E6 cells for 1 h before the virus suspension was removed and different dilutions of aqueous infusions of perilla and sage were added (Fig. 1a). As control, we included coriander, a herb that does not belong to the family of *Lamiaceae* and that is to our knowledge not commonly used as medicinal herb. We applied aqueous infusions that were prepared by boiling up the coriander, perilla, and sage leaves and subsequent simmering at 60°C for 2 h to ensure efficient extraction of water-soluble components. Intriguingly, the short-term treatment with perilla and sage infusions was sufficient to significantly inhibit the replication of SARS-CoV-2 (Fig. 1b, upper panel; for details concerning the calculation of the infectivity see Methods section and Supplementary Fig. 1). This effect did not appear to constitute a general antiviral activity of the infusions, since inhibition of HSV-1 replication in treated Vero E6 cells was not observed (Fig. 1b, lower panel). To visualize the impact of the herbal teas on the SARS-CoV-2 replication, we repeated the experiment with two different doses of virus and stained the infected cells for immunofluorescence microscopy. As depicted in Fig. 1c, the number of infected cells (green) was clearly diminished after treatment with perilla and sage infusions. The antiviral activity was still evident even when a high amount of virus was used for infection (0.5 PFU/cell). When we evaluated different members of the family of *Perilla frutescens* (red perilla, green perilla, and bi-color perilla), we observed antiviral activity in all three cases (Fig. 1d). No comparable antiviral activity was observed under the aforementioned conditions when ginger, fennel, chamomile flower, and Greek mountain teas were tested (data not shown). To confirm, that the experimental setup allows reporting of antiviral activity affecting post-entry steps, we included a SARS-CoV-2 convalescent serum sample with shown neutralizing capacity ^29^ in our analysis. We observed that the perilla and sage infusions outperformed the effect of the convalescent serum (Fig. 1e; NAbs, neutralizing antibodies) under these experimental conditions, showing that the herbs perilla and sage contain components active against SARS-CoV-2 replication by interfering with a post-entry step. Since the components were extracted by boiling the herbs in water, we concluded that the antiviral activity is elicited by water-soluble heat-stable compound(s). When we used a second clinical SARS-CoV-2 isolate to test the susceptibility towards the herbal components, we observed almost identical dose responses (Supplementary Fig. 2). The combined results of several independent experiments (using two distinct SARS-CoV-2 isolates, B.1 and B.1.1.232, for the infection of Vero E6 and α-S or α-N antibodies for staining) revealed highly significant antiviral activity of all tested dilutions of the perilla infusion as well as of the 1/10 and 1/20 dilutions of the sage infusion (Supplementary Fig. 2). Cell viability was determined to exclude cytotoxicity as reason for diminished viral replication (Supplementary Fig. 3a). In conclusion, the *Lamiaceae* perilla and sage contain water-soluble heat-stable components that exhibit potent antiviral activity against SARS-CoV-2 *in vitro*.

**Fig. 1:**
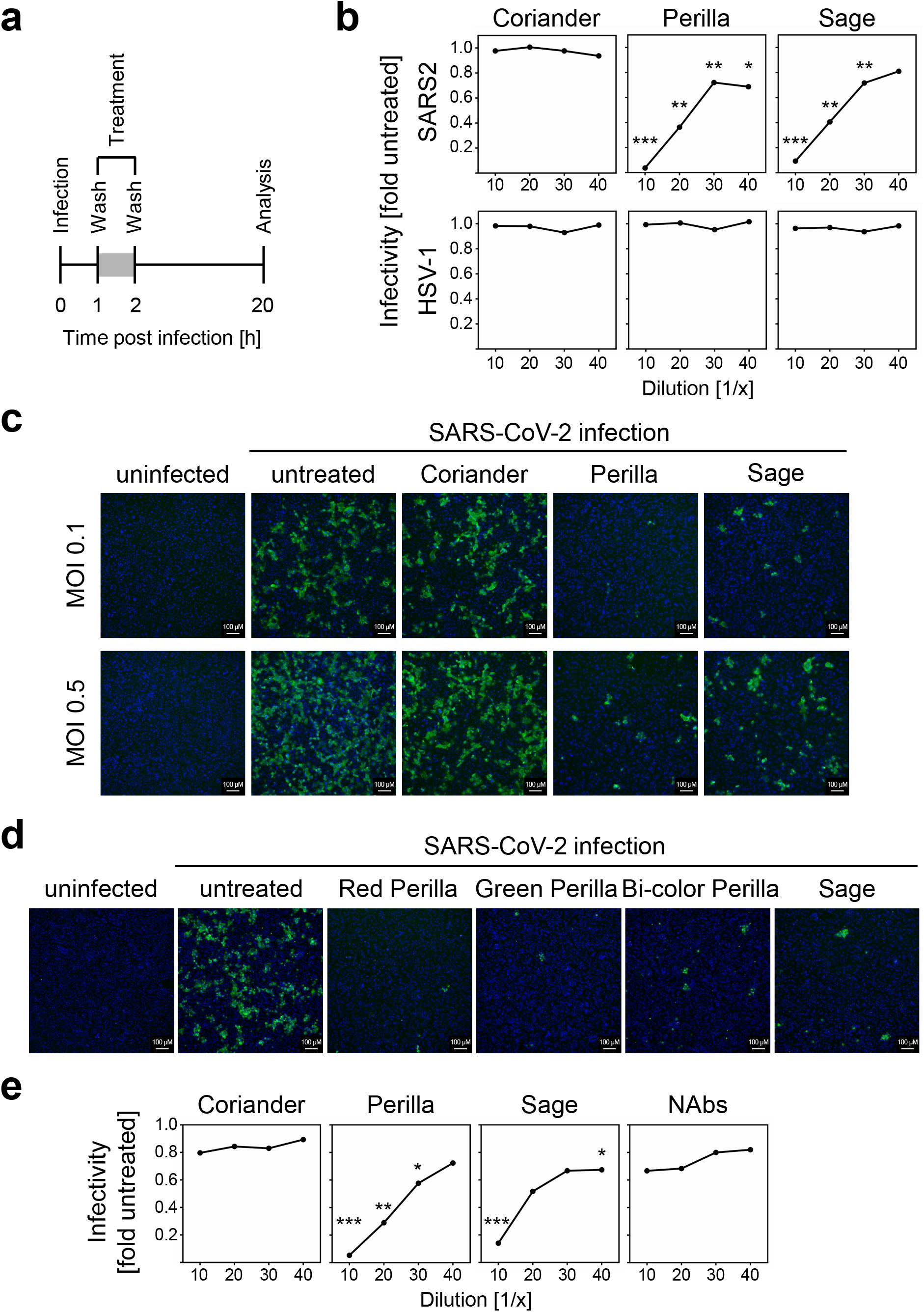
Perilla and sage contain water-soluble heat-stable components active against SARS-CoV-2 *in vitro* replication. **a** Scheme of the experimental setup for the *in vitro* analysis of therapeutic effects against SARS-CoV-2. **b** Representative dose-response curves of SARS-CoV-2-infected Vero E6 cells (2000 PFU per well) after treatment with aqueous infusions of coriander, perilla, or sage. Upper panel shows the effect on SARS-CoV-2 replication. Lower panel depicts the effect on HSV-1 replication. SARS-CoV-2 replication was analyzed at 20 h p.i. by icELISA, HSV-1:GFP replication was determined at 48 h p.i. by quantification of fluorescence. Data are expressed as relative change in infectivity compared to the untreated control. Each condition was analyzed in triplicate. See methods section and Supplementary Fig. 1 for details. The comparison of the herb-treated samples of SARS-CoV-2 to the untreated controls by one-way ANOVA showed for all dilutions of coriander no significance and for all dilutions of perilla and sage significance. The perilla- and sage-treated conditions of SARS-CoV-2 were also compared to the corresponding coriander-treated condition (same dilution) and these results are depicted in the diagram. *, p<0.05. **, p<0.01. ***, p<0.001. The comparison of the herb-treated samples of HSV-1 to the untreated controls showed for all dilutions of all tested herbs no significance. **c, d** Visualization of SARS-CoV-2 infection upon treatment with herbal infusions. Vero E6 cells were infected (MOI 0.1) and treated (1/10 dilution) as shown in **a**. α-S mAb and a Cy2-coupled secondary antibody were used for immunofluorescence staining (green). Nuclei were counterstained with DAPI (blue). **e** Representative dose-response curves of SARS-CoV-2-infected Vero E6 cells (2000 PFU per well) after treatment (as shown in **a**) with aqueous infusions of coriander, perilla, sage, or SARS-CoV-2 convalescent serum (serum 6 from ^29^ with mid-high 50% neutralization titer of 256). Each condition was analyzed in triplicate. The perilla- and sage-treated conditions were compared to the corresponding NAbs-treated condition (same dilution) by one-way ANOVA. **, p<0.01. ***, p<0.001.

### Preserved sage and perilla leaves retain bioactive compounds

Considering that the herbal infusions were prepared from 150 g of fresh herbal material per liter, which correspond to 15-30 g of dried herbal material per liter (assuming a water content of 80-90% of fresh herbs), a 1/10 dilution of these infusions already meets the concentration of herbal teas prepared from standard tea bags (1.5-2.25 g per cup). This indicates that herbal teas prepared from commercially available tea bags or dried herb leaves might contain a sufficient concentration of the antiviral compound(s). To further adapt the experimental setup to realistic conditions of tea consumption, we shortened the treatment time from 1 h to 30 min. Even under this condition, significant reduction of viral replication was observed (Supplementary Fig. 3b). Next, we compared the antiviral activity of infusions prepared from fresh or dried herb leaves. As shown in Supplementary Fig. 3c, dried sage leaves retained most of the antiviral component(s) whereas dried perilla was less effective as compared to fresh perilla leaves although significant inhibition of viral replication was still observed for the 1/10 dilutions. To assess whether conservation of perilla by freezing could be superior to drying in terms of preserving the antiviral component(s), we first tested if the antiviral activity of perilla infusions is reduced by freeze-thaw cycles. Since this did not seem to be the case (Supplementary Fig. 3d), infusions prepared from fresh and frozen perilla leaves were compared. The comparison revealed that the preservation of the herbs by freezing was preferable to drying (Supplementary Fig. 3e). Since the herbal infusions were prepared by boiling, simmering, and over-night incubation (see Methods section), we also tested if the standard procedure of herbal tea preparation using dried sage leaves is sufficient to extract the antiviral component(s). To this end, dried sage leaves were boiled up in water and incubated for 10 min before the herb leaves were removed. When this 10-min infusion of dried sage was compared to the over-night infusion, very similar dose response curves were observed (Supplementary Fig. 3f).

### Perilla and sage elicit prophylactic antiviral activity *in vitro*

Having observed the potent antiviral activity of perilla and sage after only 1 h of treatment, we wondered whether the herbs might also elicit prophylactic effects. Therefore, Vero E6 cells were treated 1 h prior to infection with different dilutions of the herbal infusions before the supernatant including the herbal components was removed. Subsequently, SARS-CoV-2 infection was performed and the virus suspension was replaced by fresh medium at 1 h p.i. (Supplementary Fig. 4a). By removing the herbal infusions before infection, we aimed to primarily assess antiviral effects based on cellular responses and not on direct virucidal elimination of infectious virus particles. The analysis of combined results of 6 independent experiments using two distinct SARS-CoV-2 isolates and α-S or α-N antibodies for staining showed highly significant decrease of infectivity, especially upon pre-treatment with the perilla infusion (Supplementary Fig. 4b). To compare the extent of prophylactic and therapeutic antiviral capacity, we conducted an experiment in which we treated and pre-treated the infected cells in parallel. As already indicated by the results of the independent experiments (Supplementary Fig. 2 a, b and Supplementary Fig. 4b), therapeutic treatment elicited stronger antiviral activity for both perilla and sage (Supplementary Fig. 4 c, d). Nonetheless, perilla and sage teas might not only be suitable for treatment of SARS-CoV-2 infections but also for prevention of infections.

### Perilla and sage confer protection against SARS-CoV-2 infection in human cells

Since cells differ concerning the mode of entry of SARS-CoV-2 ^30^, a second independent cell line was tested. Caco-2 cells were used to analyze the antiviral activity elicited by perilla and sage infusions in human cells. We have observed that SARS-CoV-2 replication is more protracted in Caco-2 cells compared to Vero E6 cells ^29^. Therefore, the experimental setup was adapted by increasing the time of treatment as well as the time of infection before analysis (Supplementary Fig. 5a). We observed a strong decrease in infectivity in perilla- and sage-treated Caco-2 cells (Supplementary Fig. 5b). Encouraged by this result, we applied the same treatment regimen of 1 h as was used for Vero E6 cells (Fig. 2a). This early short-term treatment was sufficient to potently inhibit SARS-CoV-2 replication, even in cells with a protracted viral replication cycle (Fig. 2b). To visualize the antiviral activity, Caco-2 cells were infected with and without treatment (as depicted in Fig. 2a) and were fixed for fluorescence microscopic analysis. This analysis showed clearly visible differences in the number of Spike-positive cells (Fig. 2c). To corroborate the data obtained so far, we analyzed the antiviral activity of herbal infusions by staining the intracellular spike protein by in-cell-ELISA (icELISA ^29^) as well as quantifying the viral genomes in the supernatant by qRT-PCR. The results showed that the icELISA data reflected the decrease in viral replication and the resulting decline in viral progeny (Fig. 2d).

**Fig. 2:**
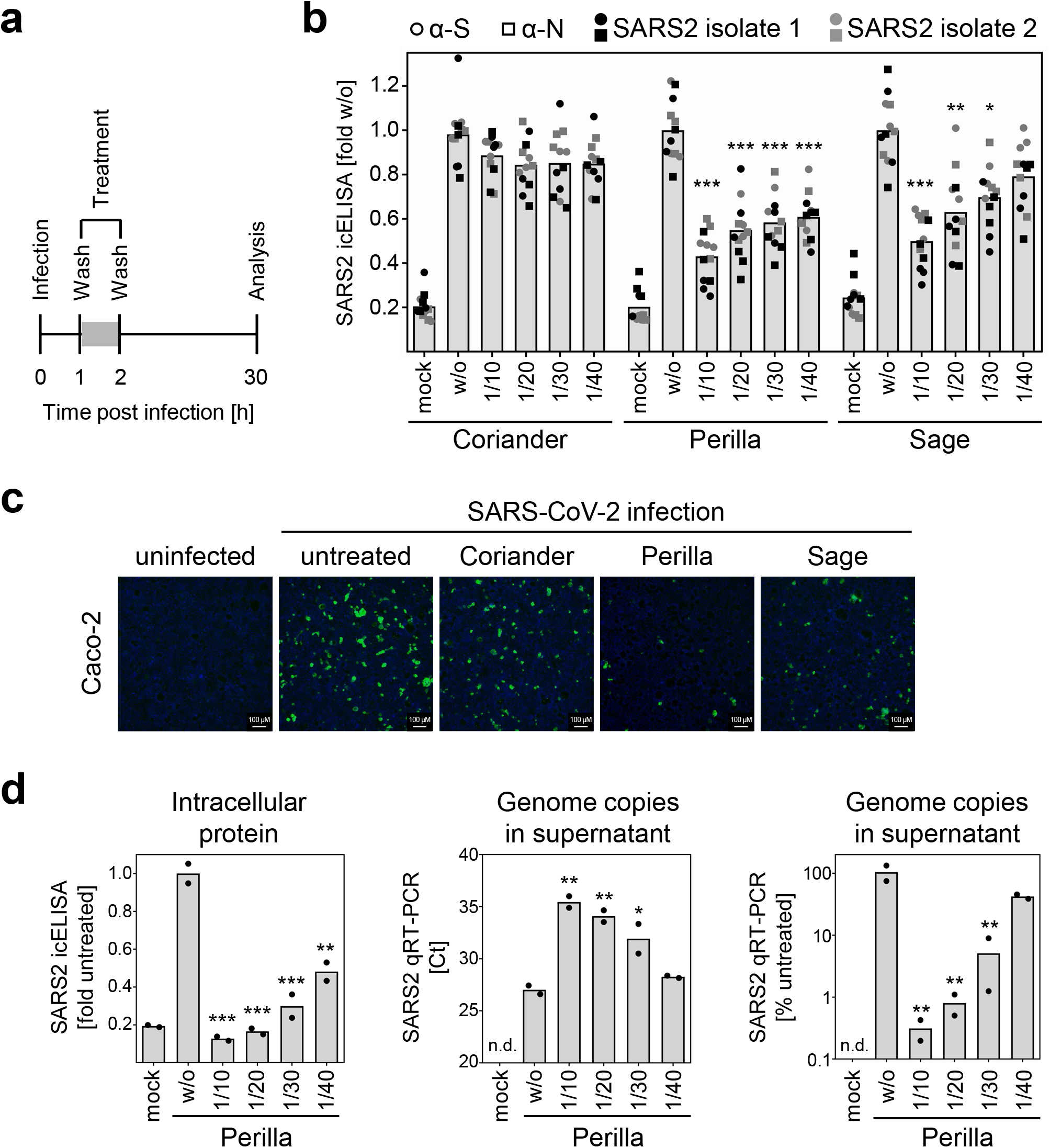
Perilla and sage confer protection against SARS-CoV-2 infection in human cells. **a** Scheme of the experimental setup for the *in vitro* analysis of antiviral activity against SARS-CoV-2 in human Caco-2 cells. **b** Pooled icELISA data of 4 independent experiments using two distinct SARS-CoV-2 isolates (B.1. and B1.1.232) for infection of Caco-2 cells and α-S or α-N mAbs for staining. Data are expressed as relative change in optical density compared to the untreated control. The perilla- and sage-treated conditions were compared to the corresponding coriander-treated condition (same dilution) by one-way ANOVA. *, p<0.05. **, p<0.01. ***, p<0.001. **c** Visualization of SARS-CoV-2 infection upon treatment with herbal infusions. Human Caco-2 cells were infected and treated as shown in **a**. α-S mAb and a Cy2-coupled secondary antibody were used for immunofluorescence staining (green). Nuclei were counterstained with DAPI (blue). **d** Human Caco-2 cells were infected and treated as shown in Supplementary Fig 5 **a**. At 30 h p.i., supernatant was collected for RNA preparation and subsequent qRT-PCR analysis. Cells were fixed and analyzed by icELISA using α-S mAb for staining. icELISA data are expressed as relative change in optical density compared to the untreated control. qRT-PCR data are shown in Ct value and calculated relative change in genome copies compared to the untreated control. Each condition was analyzed in duplicate. The perilla-treated conditions were compared to the untreated control by one-way ANOVA. *, p<0.05. **, p<0.01. ***, p<0.001.

### The herbal infusions exert antiviral effects comparable to IFNβ and remdesivir and outperform IFNα2 upon short-term treatment

Interferons mediate innate protection against viruses and are considered a treatment option for COVID-19 disease and provide certain benchmark for relevant antiviral activity. Another antiviral drug authorized for emergency use in COVID-19 patients is remdesivir, an adenosine nucleotide analog prodrug inhibiting the viral RNA-dependent RNA polymerase. To assess the potency of the herbal infusions, we compared treatment with perilla and sage infusions with treatment with remdesivir, IFNα2, and IFNβ. We infected Vero E6 or Caco-2 cells with SARS-CoV-2 and treated the infected cells at 1 h post-infection for 1 h before the treatment was removed. The antiviral activity was determined by quantification of viral proteins by icELISA. Under herein used experimental condition, perilla and sage infusions showed higher antiviral potency than remdesivir in Vero E6 cells (Fig. 3a) and comparable antiviral activity in infected Caco-2 cells (Fig. 3b). When the herbal infusions were compared to IFNs, we observed similar effects to short-term IFNβ treatment but higher potency than short-term treatment with IFNα2 (Fig. 3c). Consistent with the notion that the herbal infusions act in a different manner than the IFNs, treatment with the Janus kinase inhibitor ruxolitinib abrogated the antiviral activity of IFNβ but not perilla and sage (Fig. 3d).

**Fig. 3:**
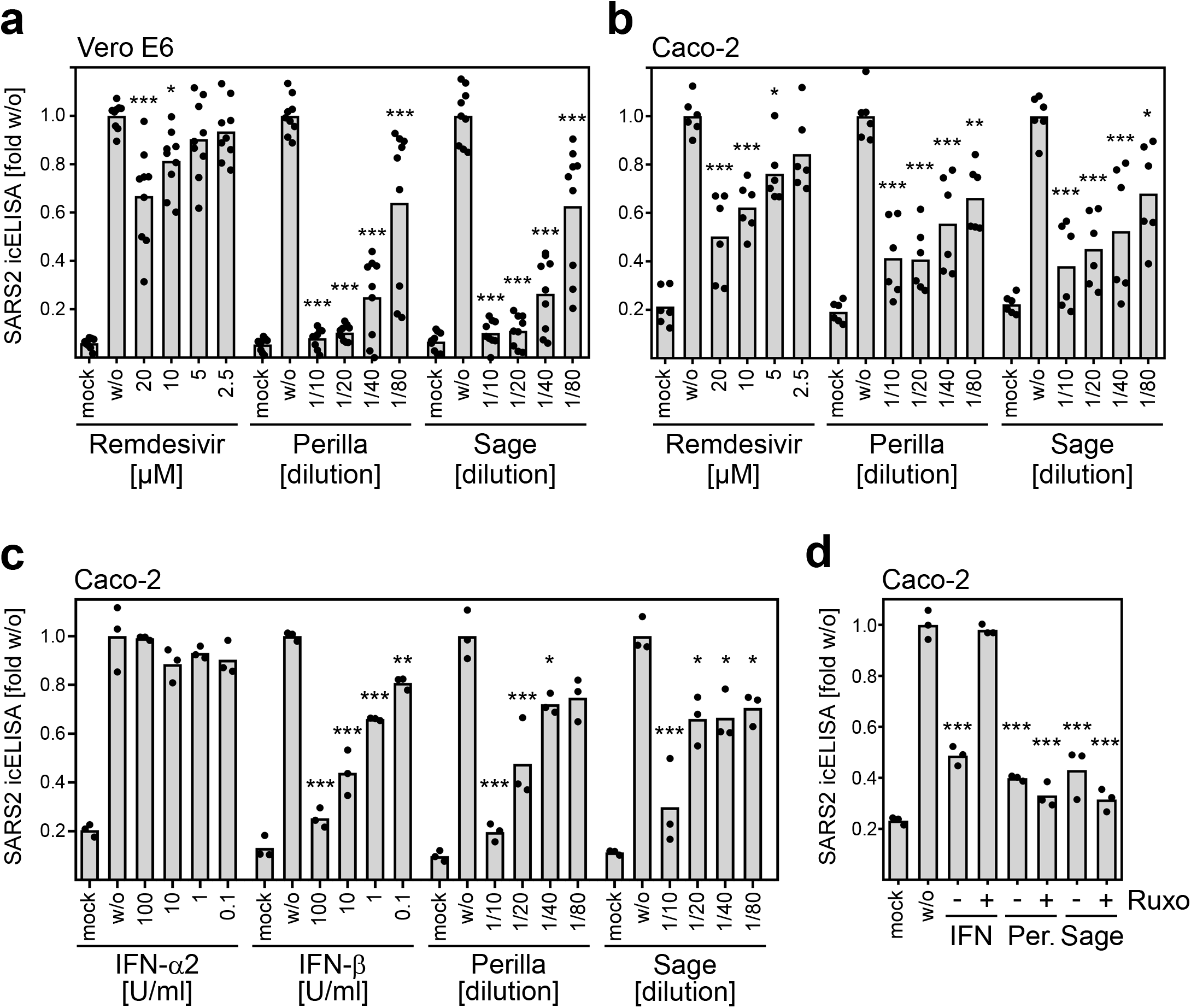
The herbal infusions exerted comparable antiviral effects to IFNβ and remdesivir and outperformed IINα2 upon short-term treatment. **a, b** Pooled icELISA data of SARS-CoV-2-infected cells (**a**, Vero E6, 3 independent experiments; **b**, Caco-2, 2 independent experiments) after treatment with herbal infusions or remdesivir at 1 h p.i. for 1 h. Vero E6 and Caco-2 were fixed at 24 h p.i. and stained with α-S and α-N, respectively. Data are expressed as relative change in optical density compared to the untreated control. The treated conditions were compared to the untreated control by one-way ANOVA. *, p<0.05. **, p<0.01. ***, p<0.001. **c** icELISA data of SARS-CoV-2-infected Caco-2 cells after treatment with herbal infusions or indicated interferons at 1 h p.i. for 1.5 h. Cells were fixed at 30 h p.i. and stained with α-S mAb. Each condition was analyzed in triplicate. The treated conditions were compared to the untreated control by one-way ANOVA. *, p<0.05. **, p<0.01. ***, p<0.001. **d** icELISA data of SARS-CoV-2-infected Caco-2 cells after treatment with IFNβ or herbal infusions (P, perilla; S, sage) at 1 h p.i. for 1.5 h in the absence or presence of the Janus kinase inhibitor ruxolitinib (4 μM). The treated conditions were compared to the untreated control by one-way ANOVA. ***, p<0.001.

### Identification of antiviral components following size exclusion and protein fractionation analyses

Since the herbal infusions contain a large number of different components, it appeared difficult to determine the active substances without obtaining additional information about the nature of these active components. To narrow down candidate substances, we fractionated the components of the herbal infusions according to their molecular size and tested the antiviral activity of these fractions. When we tested the fractions obtained by use of Amicon 100K, 30K, and 10K filters, we were able to assign the active component(s) to substances less than 10 kDa in size (Supplementary Fig. 6). To further restrict the number of substances to be analyzed, we used Amicon 10K and 3K filters for fractionation. Additionally, we prepared by dialysis the fraction of substances larger than 1 kDa. The analysis of these fractions revealed that the dialyzed herbal infusions exhibited decreased antiviral activity indicating that components less than 1 kDa in size contribute to the antiviral activity of perilla (Fig. 4a) and sage infusions (Fig. 4b). In light of these findings, we reviewed the literature for candidate substances. Based on published lists of perilla components (e.g., ^31,32^), we tested various compounds present in *Lamiaceae*. We did not detect antiviral activity against SARS-CoV-2 when we tested apigenin, luteolin, rosmarinic acid, and tormentic acid (data not shown). Interestingly, when we analyzed cinnamic acid, hydroxy-cinnamic acid, and dihydroxy-cinnamic acid, only the treatment with dihydroxy-cinnamic acid diminished SARS-CoV-2 replication in Vero E6 (data not shown) and Caco-2 cells (Supplementary Fig. 7a, Fig. 4c). This finding is remarkable, since hydroxy-cinnamic acid and dihydroxy-cinnamic acid (that is more familiar under the name caffeic acid) differ only by a single hydroxyl group. In addition, we found that, besides caffeic acid, perilla aldehyde and perillyl alcohol also exhibit antiviral activity against SARS-CoV-2 (Supplementary Fig. 7b, Fig. 4c). These results suggest that the effect of the herbal infusions is based on multiple components that presumably act through additive mechanisms. To evaluate whether knowledge of the active components could help predict additional antiviral herbs, we searched for herbs containing high amounts of caffeic acid. Thyme is a herb with a high content of caffeic acid (http://phenol-explorer.eu/contents/polyphenol/457). Therefore, we analyzed aqueous infusions of thyme. We observed potent antiviral activity against SARS-CoV-2 in Vero E6 and Caco-2 cells (Supplementary Fig. 7c). Thus, the identification of antiviral herbal components enabled the prediction of further antiviral herbs and may pave the way to novel therapeutics for COVID-19.

**Fig. 4:**
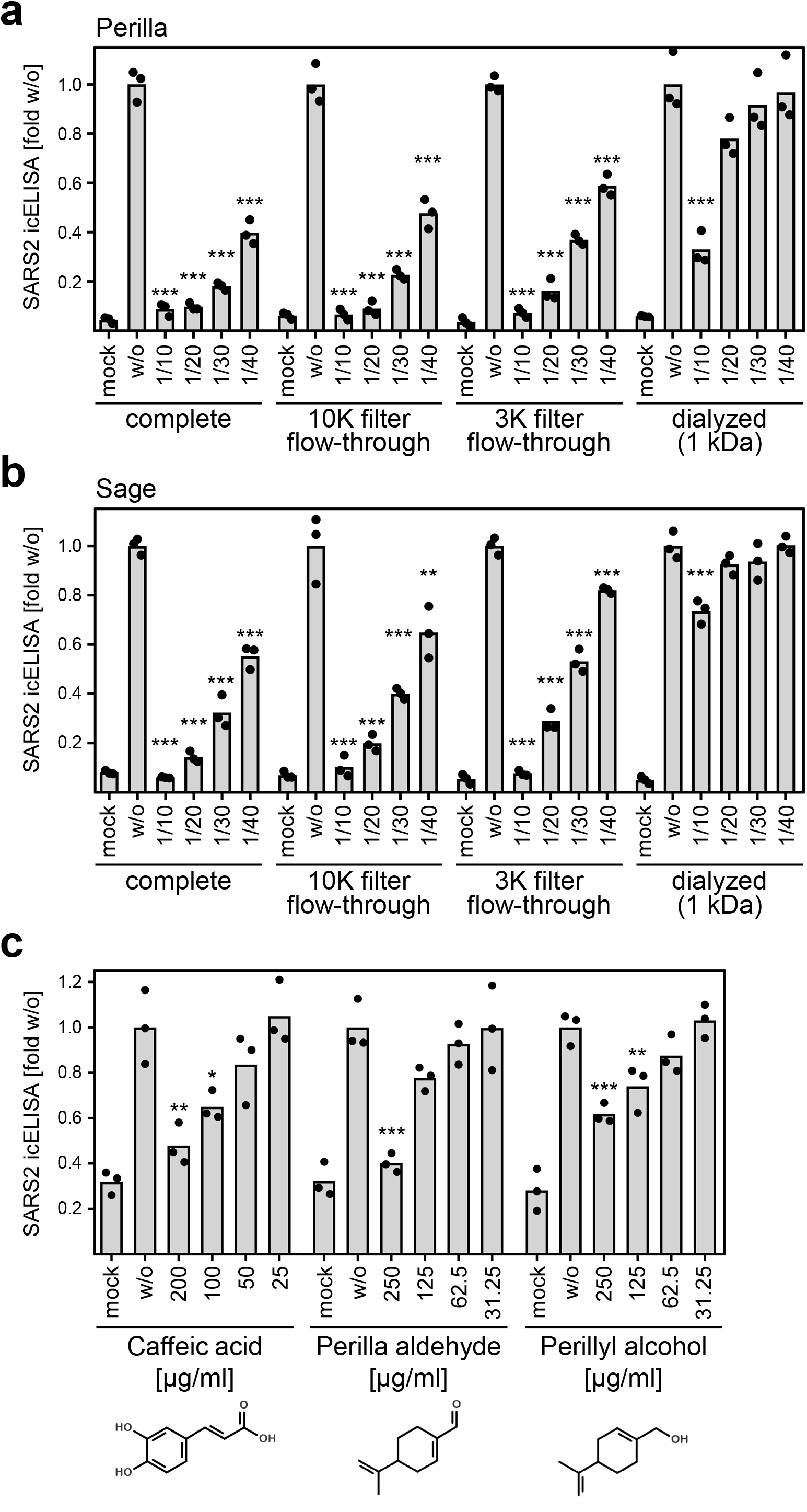
Identification of antiviral components following size exclusion and protein fractionation analyses. **a, b** icELISA data of SARS-CoV-2-infected Vero E6 cells after treatment with indicated fractions of herbal infusions (**a**, perilla; **b**, sage) at 1 h p.i. for 1 h. Cells were fixed at 20 h p.i. and stained with α-S mAb. Data are expressed as relative change in optical density compared to the untreated control. Each condition was analyzed in triplicate. Protein fractionation of herbal infusion components was conducted by use of Amicon 10K and 3K filters. The fraction of proteins >1 kDa was obtained by dialysis. Dilutions and concentrations by the fractionation steps were considered. The treated conditions were compared to the untreated control by one-way ANOVA. **, p<0.01. ***, p<0.001. **c** icELISA data of SARS-CoV-2-infected Caco-2 cells after treatment with components of herbal infusions at 1.5 h p.i. for 1.5 h. Cells were fixed at 24 h p.i. and stained with α-N mAb. Each condition was analyzed in triplicate. The treated conditions were compared to the untreated control by one-way ANOVA. *, p<0.05. **, p<0.01. ***, p<0.001.

### Global mass spectrometry (MS) analyses proposed induction of HMOX-1 as anti-SARS-CoV-2 effector mechanism of herbal infusions

Next, we sought to determine proteomic changes in cells treated with perilla and sage infusions to advance our understanding of effector mechanisms that lead to the reduction of SARS-CoV-2 replication. To this end, SARS-CoV-2-infected Vero E6 and Caco-2 cells as well as mock controls were treated at 1 h p.i. with perilla and sage infusion for 1 h. At different time points after infection (2, 6, 20 h p.i. in the case of Vero E6 and 2, 6, 30 h p.i. in the case of Caco-2), cells were lysed and subjected to global MS analysis. To be able to detect the mutual interplay between viral and host proteins, we increased the virus amount used for infection to MOI 0.3 and MOI 0.5 for Vero E6 and Caco-2, respectively. This experimental setting might result in overwhelming expression of viral gene products at late times post-infection, but it presumably allows sufficient viral gene expression early after infection to unravel the impact of viral proteins on cellular processes. Overall, we detected and quantified 19 viral proteins and protein forms as well as 3733 cellular proteins in Vero E6 and 4646 cellular proteins in Caco-2 samples by MS. To our knowledge, this is the first reported proteomic dataset investigating effects of herbal extracts such as perilla and sage in cell culture models. To support broad access to the dataset and to allow facile data visualization, we included a ready-to-use Microsoft Excel-based plotting option as supplement (Supplementary File 1 and 2 for quantified Vero E6 and Caco-2 proteins, respectively). Using these tables, relative abundances of all individual proteins detected by at least two unique peptides can be visualized. Next, we examined the MS data to evaluate whether proteomic changes in host proteins, which have been suggested as host restriction factors (HRFs) and host dependency factors (HDFs) ^33-35^, could explain the observed antiviral activity. For this purpose, we analyzed all proposed HRFs and HDFs that could be consistently quantified across all investigated conditions. These analyses revealed no significant and consistent upregulation of HRFs and downregulation of HDFs upon treatment with perilla or sage infusions (Supplementary Fig. 8 and 9). Therefore, we searched for proteins significantly regulated at least 1.5-fold upon treatment with perilla or sage infusions in infected cells at 6 h p. i. because this condition most likely includes the relevant changes. For the differential comparisons, the proteins were filtered by the coefficient of variation (CV): all proteins with a CV <20% over all measurements (also across treatments) were not considered. When comparing the changes induced by treatment with perilla or sage infusions in infected cells at 6 h p.i., Heme Oxygenase 1 (HMOX-1) was the only protein found to be significantly altered under all 4 conditions (Fig. 5a). HMOX-1 is an enzyme involved in the response to oxidative stress. It catalyzes the oxidative degradation of heme to biliverdin, a precursor of bilirubin, thereby detoxifying free heme ^36,37^. When we examined in detail the individual proteins regulated by perilla and sage infusions, we found additional proteins involved in the oxidative stress response (Fig. 5b and Supplementary Fig. 10). As example, Sulfiredoxin 1 (SRXN1), an endogenous antioxidant protein that prevents cell oxidative stress damage, was also upregulated in Caco-2 cells after treatment. Interestingly, levels of CYP1A1, that was recently shown to inhibit HMOX-1-mediated bilirubin formation ^38^, were decreased in infected Caco-2 cells after treatment with perilla or sage infusions (Fig. 5b and Supplementary Fig. 10). This finding confirmed that activity of HMOX-1 was elevated in infected cells treated with perilla or sage infusions. Noteworthy, viral N as well as membrane (M) and ORF7a (7a) proteins were the most strongly downregulated proteins in Caco-2 and Vero E6 cells, respectively, when treated with herbal infusions (Fig. 5b and Supplementary Fig. 10), which further corroborates the observed antiviral activity.

**Fig. 5:**
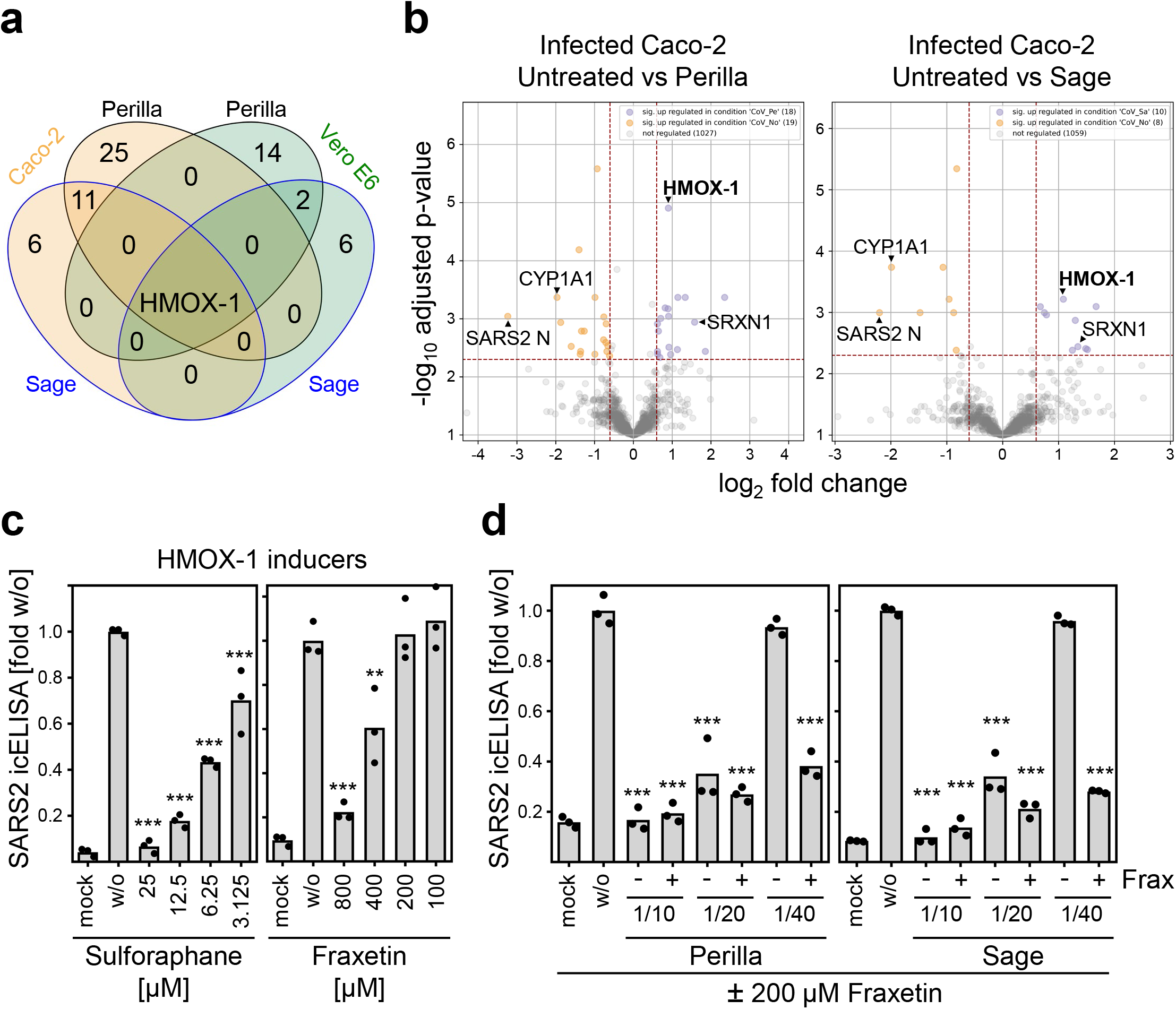
Global MS analyses proposed HMOX-1 induction as anti-SARS-CoV-2 effector mechanism. **a** SARS-CoV-2-infected Vero E6 and Caco-2 cells were treated at 1 h p.i. with perilla or sage infusions for 1 h. At 6 h p.i., cells were lysed and subjected to global MS analysis (see methods section for details). Untreated and infusion-treated cells infected with SARS-CoV-2 were compared (see methods section for details), overlapping significantly regulated proteins are shown. **b** Volcano plots of MS results obtained from infected Caco-2 at 6 h p.i. Normalized data were filtered (at least two unique peptides per protein group required) and proteins with a coefficient of variation <20.0% over all measurements (also across treatments) were removed (see methods section for details). **c** SARS-CoV-2-infected Vero E6 cells were treated at 1 h p.i. for 1.5 h with the HMOX-1 inducer sulforaphane or fraxetin. Cells were fixed at 24 h p.i. and stained with α-S mAb. Data are expressed as relative change in optical density compared to the untreated control. Each condition was analyzed in triplicate. The treated conditions were compared to the untreated control by one-way ANOVA. **, p<0.01. ***, p<0.001. **d** SARS-CoV-2-infected Vero E6 cells were treated at 1 h p.i. for 1.5 h with 200 μM fraxetin and indicated dilutions of perilla or sage infusions. At 24 h p.i., cells were fixed and stained with α-S mAb. Data are expressed as relative change in optical density compared to the untreated control. Each condition was analyzed in triplicate. The treated conditions were compared to the untreated control by one-way ANOVA. ***, p<0.001.

To test whether the increase in HMOX-1 protein levels and activity may represent one of the effector mechanisms of herbal infusions to restrict SARS-CoV-2 replication, we examined the antiviral activity of HMOX-1-inducing substances such as sulforaphane and fraxetin. Indeed, we observed antiviral activity after short-term treatment with sulforaphane or fraxetin (Fig. 5c). This result raised the question of whether this information could be applied to improve the efficacy of the herbal treatments. HMOX-1 was found to be upregulated about 2-fold in infected Caco-2 cells (Fig. 5b). Therefore, we wondered whether increased HMOX-1 induction could boost the antiviral activity of perilla and sage. To test this option, we combined the herbal infusions with a dose of fraxetin that was in itself not active against SARS-CoV-2 to treat infected cells. In fact, the combination of low-dose fraxetin with perilla or sage infusions strongly enhanced the antiviral activity (Fig. 5d).

### Herbal infusions exhibit antiviral activity against SARS-CoV-2 variants of concern

Since the beginning of the pandemic, several SARS-CoV-2 variants emerged. In particular, the variants of concern (VOCs) exhibit strong impact on the course of the pandemic. Therefore, we tested whether the antiviral principle induced by treatment with perilla and sage infusions also counteracts infections with the VOCs Alpha (B.1.1.7), Beta (B.1.351), Delta (B.1.617.2), and Omicron (BA.1). The E484K mutation present in the S protein of the Beta variant was shown to be an escape mutation impairing the neutralizing capacity of antibodies ^39,40^ This finding was confirmed with our Beta virus isolate. We were not able to stain Beta by icELISA when we used two different monoclonal antibodies both binding to the RBD of S (Supplementary Fig. 11a). When using a polyclonal α-RBD antibody, the signal was reduced but not completely lost (Supplementary Fig. 11a). In contrast, the recognition by α-N antibodies was not affected (Supplementary Fig. 11a). Based on these observations, we used α-N for icELISA detection and quantification of Beta infections. We analyzed the antiviral activity of perilla and sage infusions by icELISA as well as qRT-PCR quantification of viral progeny in the supernatant and observed similar antiviral effects against wild-type, Alpha, and Beta (Supplementary Fig. 11b, Fig. 6a). We also detected strong antiviral activity of perilla and sage infusions against the Delta variant (Fig. 6b), but not when coriander infusion was used (data not shown). Delta could only be tested in human cells because the P681R mutation in the S protein of Delta almost completely prevents infection of Vero E6 cells (unpublished observation). Similarly, the currently most widespread Omicron variant was assessed in human cells. In our hands, Omicron hardly replicated in Vero E6 cells, and even in human cells, Omicron replicated only to low titers (unpublished observation). For this reason, we used lower input virus doses and did not apply the icELISA readout but used the more sensitive quantification of viral progeny by qRT-PCR. This analysis revealed that perilla and sage infusions also elicited antiviral activity against Omicron (Fig. 6c).

**Fig. 6:**
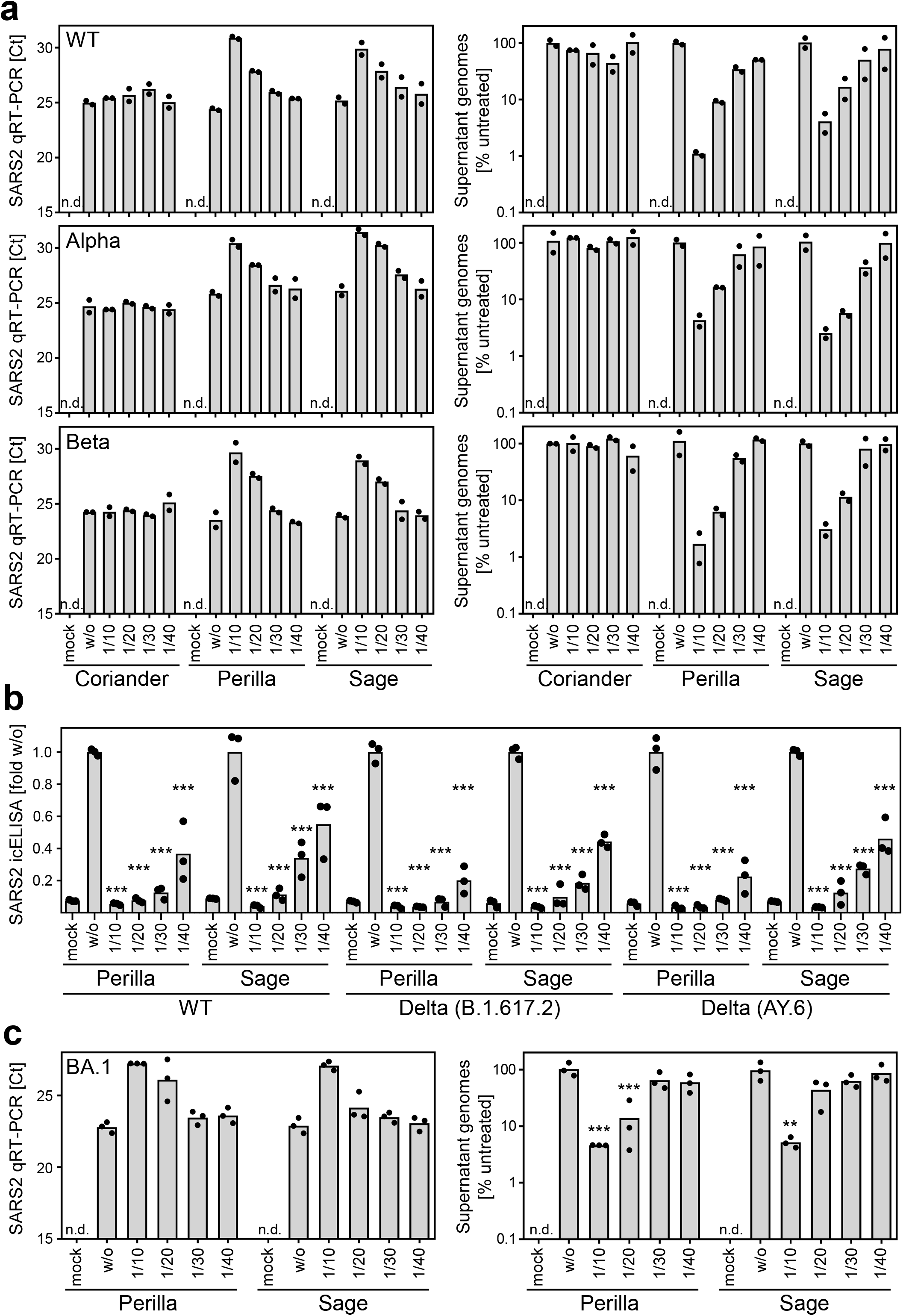
Herbal infusions exhibit antiviral activity against SARS-CoV-2 variants of concern. **a** Vero E6 cells were infected with SARS-CoV-2 wt and the variants of concern Alpha (B.1.1.7) and Beta (B.1.351) after treatment with aqueous infusions of coriander, perilla, or sage at 1 h p.i. for 1 h. SARS-CoV-2 replication was determined by quantification of viral genomes in the supernatant of infected cells at 20 h p.i.. Data are shown in Ct value and calculated relative change in genome copies compared to the untreated control. Each condition was analyzed in duplicate. **b** Caco-2 cells were infected with SARS-CoV-2 wt and two distinct isolates of Delta (B.1.617.2 and AY.6) after treatment with aqueous infusions of perilla or sage at 1.5 h p.i. for 1.5 h. SARS-CoV-2 replication was analyzed at 30 h p.i. by icELISA (α-S staining). Data are expressed as relative change compared to the untreated control. Each condition was analyzed in triplicate. The treated conditions were compared to the untreated control by one-way ANOVA. ***, p<0.001. **c** Caco-2 cells were infected with SARS-CoV-2 variant of concern Omicron (BA.1) after treatment with aqueous infusions of perilla or sage at 1.5 h p.i. for 1.5 h. SARS-CoV-2 replication was determined by quantification of viral genomes in the supernatant of infected cells at 24 h p.i.. Data are shown in Ct value and calculated relative change in genome copies compared to the untreated control. Each condition was analyzed in triplicate. The treated conditions were compared to the untreated control by one-way ANOVA. ***, p<0.001. **, p<0.01.

Taken together, aqueous infusions based on perilla and sage exhibit strong *in vitro* antiviral activity against SARS-CoV-2 including VOCs such as Alpha, Beta, Delta, and Omicron.

## Discussion

The WHO SOLIDARITY trial comprising more than 400 hospitals and 11,000 patients in 30 countries showed that “*Remdesivir, Hydroxychloroquine, Lopinavir and Interferon regimens appeared to have little or no effect on hospitalized COVID-19, as indicated by overall mortality, initiation of ventilation and duration of hospital stay*” ^41^. To a certain extent, this finding may be attributed to the practice of reserving such drugs for critically ill patients for whom they may be applied too late in terms of disease progression. Recently, Merck and Pfizer applied for an emergency use authorization by the US Food and Drug Administration (FDA) for the drugs Molnupiravir and Paxlovid, respectively. Clinical trials showed promising results in reducing the risk of hospitalization or death when given early in the infection, but definitive clinical data are still needed to prove the efficacy of the antiviral treatment candidates. A widely discussed potential solution to the limited efficacy of single therapies is the combination of substances that have a marginal effect on their own. Such “shotgun” treatment regimens have already shown some success ^42^. Since herbal teas are ubiquitously available, almost free of charge, and exhibit excellent safety profiles given their consumption as spices, we propose to add perilla, sage, and thyme teas to combinatorial treatment regimens. Furthermore, these *Lamiaceae* teas may be applicable as an addition to the very important non-pharmaceutical interventions such as wearing a face cover, proper hygiene, physical distance, and the restriction of social interactions.

Using different SARS-CoV-2 cell culture models, highly permissive Vero E6 cells and human Caco-2 cells, we observed potent antiviral activity of herbal teas produced by boiling leaves of the *Lamiaceae* perilla and sage. A typical cup of tea corresponds to approx. 250 ml of volume and commercial tea bags usually contain 1.5-2.25 g of plant material. Our herbal extracts were produced using 15 g of fresh herb leaves boiled in 100 ml of water. Assuming a water content of 80-90% of fresh herbs, a 1/10 dilution of these infusions already corresponds to herbal teas prepared from standard tea bags (1.5-2.25 g per cup). In human cells, perilla and sage infusions elicited significant antiviral effects at dilutions of 1/40 (3.75 g per liter or ca. 0.94 g per cup) and 1/30 (5 g per liter or ca. 1.25 g per cup) (Fig. 2b), indicating efficacy at concentrations usually consumed in such herbal teas.

Given that Vero cells are incapable to express type I interferons due to genetic aberrations ^43^, the antiviral activity observed in Vero E6 cells indicate that perilla and sage extracts elicit their effect independent of interferon induction. The result that the Janus kinase inhibitor ruxolitinib did not revert the antiviral activity elicited by perilla and sage infusions in Caco-2 cells supports this conclusion. Our finding that *Lamiaceae* infusions exhibit antiviral activity in both Vero E6 and Caco-2 cells (Fig. 1 and 2) suggests that S cleavage and the endosomal entry route are at least not the only target of the antiviral mechanism, since SARS-CoV-2 entry into Vero E6 and Caco-2 is mediated by distinct routes ^30^. This conclusion is further supported by our observation that sage and perilla teas diminished viral replication, especially when they were applied after infection at a time point at which neutralizing antibodies present in convalescent serum had lost most of their antiviral activity. In such treatment regimens, the herbal teas significantly outperformed NAb-containing serum (Fig. 1e).

Natural products and traditional herbal medicines have often paved the way to the development of novel therapeutic agents. Many drugs were developed based on the structure of natural compounds that exhibit the desired effect. Examples of this approach of drug development include the antimalarial drug artemisinin derived from *Artemisia annua*, the cancer drug taxol derived from *Taxus brevifolia*, and the amoebicide emetine, an isoquinoline alkaloid from *Cephaelis ipecacuanha*. Numerous other drugs were modified from natural compounds such as aspirin and morphine. In fact, half of all drugs approved between 1981 and 2019 were derived from or mimicked a natural compound ^44^.

Upon selection of candidate compounds based on protein fractionation analyses, we identified caffeic acid, perilla aldehyde, and perillyl alcohol as antiviral substances (Fig. 4). Caffeic acid is a plant-derived compound containing both phenolic and acrylic functional groups. It is abundantly found in natural products and was shown to exhibit antimicrobial and antioxidant activities ^45,46^. The observation that caffeic acid/dihydroxy-cinnamic acid, but not hydroxy-cinnamic acid, acts antiviral (Supplementary Fig. 7a) provides insight into the reactive groups and may allow improvement of activity by structural optimization. Perilla aldehyde has gained attention in recent years because of its antioxidant and antifungal properties and its potential to serve as organic fruit and food preservative ^47^. Animal studies indicated anticancer and anti-inflammatory activity ^48^. Perillyl alcohol has also attracted attention because of its potential anticancer activity ^49^. Our data constitute the first experimental evidence for the antiviral activity of these substances against SARS-CoV-2, suggesting that caffeic acid, perilla aldehyde, and perillyl alcohol could be candidates for the development of antiviral agents that are inexpensive and therefore widely available. Plants of the *Lamiaceae* family have been shown to possess antiviral activity against the retrovirus HIV ^50-52^. Similar to our findings with SARS-CoV-2, the anti-HIV activity seems to occur at a post-entry step ^53^. It will be very interesting to test the compounds responsible for the anti-SARS-CoV-2 activity of perilla and sage infusions and to elucidate whether the antiviral activity against retroviruses and coronaviruses is mediated by the same or similar mechanisms.

Global MS analyses revealed that both perilla and sage treatments upregulated HMOX-1 abundance and activity. Follow-up experiments showed that upregulation of HMOX-1 is indeed an antiviral effector mechanism inhibiting SARS-CoV-2 infection (Fig. 5). Considering that SARS-CoV-2-induced inflammation ^54^, coagulopathies ^55^, and respiratory distress syndrome (ARDS) ^56^ are major causes of mortality in COVID-19 patients ^57,58^, the known protective role of HMOX-1 activation regarding inflammation, inflammation-induced coagulation, lung damage, and ARDS ^59-63^ suggests that herbal teas may not only help in limiting virus replication but also in treating the symptoms.

The US Food and Drug Administration (FDA) generally recognizes sage as safe ^64^, however, herbs in general and sage in particular should not be consumed by pregnant and breastfeeding women without consultation of their health-care provider ^65^ due to the potential to cause toxicity to the fetus or to induce premature labor. Sage is also discussed to reduce the milk supply when consumed during lactation. Additionally, sage and perilla can cause allergic reactions ^66^. Obviously, the consumption of herbal teas cannot and should not replace non-pharmaceutical interventions or clinically approved drugs. However, given their inexpensive and universal availability, they might contribute to prevent and/or relieve some of the hardness and suffering of the COVID-19 pandemic. Even after the pandemic phase, antiviral herbal teas could remain beneficial during endemic outbreaks. We are convinced that our data argue in favor of future clinical studies addressing the question of whether herbal teas based on perilla or sage may either be able to prophylactically reduce infections and/or offer therapeutic benefits when administered concomitantly with the standard treatment. In addition, our study identified caffeic acid, perilla aldehyde, and perillyl alcohol as potential antiviral drugs that deserve further investigations.

## Methods

### Materials and correspondence

Further information and requests for resources and reagents should be directed to and will be fulfilled by Mirko Trilling.

### Cells, viruses, and infection

Vero E6 (ATCC CRL-1586) and Caco-2 (ATCC HTB-37) were grown in high glucose Dulbecco’s minimal essential medium (DMEM [Gibco 41966-029]) and Roswell Park Memorial Institute 1640 (RPMI-1640 [Gibco 21875-034]), respectively, supplemented with 10% (v/v) FCS, penicillin, and streptomycin. Calu-3 (ATCC HTB-55) were cultivated in minimal essential medium (MEM [Gibco 31095-029]) supplemented with 10% (v/v) FCS, 1 mM sodium pyruvate, penicillin, and streptomycin. All cells were kept at 37°C in an atmosphere of 5% CO_2_.

The SARS-CoV-2 strains B.1, B1.1.232, Alpha (B.1.1.7), Beta (B.1.351), Delta (B.1.617.2 and AY.6), and Omicron (BA.1) were isolated from patient samples obtained in May 2020, March to May 2021, and December 2021. For the isolation, permissive cells were incubated with virus-containing clinical nasopharyngeal swab samples until CPE was observed. All strains except Delta and Omicron were amplified in Vero E6 cells. The Delta isolates B.1.617.2 and AY.6 as well as the Omicron isolate BA.1 were isolated and amplified using Calu-3 cells. All SARS-CoV-2 isolates were analyzed by Next Generation Sequencing (data not shown) and classified with the help of GISAID and assigned into clades according to pangolin ^67-70^. The authors made use of the outbreak.info ^71^. Viral titers were determined by 50% tissue culture infectious dose (TCID50) titration. The virus isolation has been approved by the ethics committee of the medical faculty of the University of Duisburg-Essen (20-9511-BO and 20-9512-BO). HSV-1-ΔgE-GFP was generated and described ^72^ by the laboratory of Prof. David C Johnson (Oregon Health & Science University, USA). With Prof. Johnson’s written permission, we received the virus from Prof. Hartmut Hengel (University of Freiburg, Germany).

### Generation of aqueous infusions of herbs

The herbal infusions were prepared by boiling up 15 g of fresh herbal leaves in 100 ml of water and subsequent simmering at 60°C for 2 h. The infusions were stored over-night at 4°C before the leaves were removed and the aqueous solutions were sterile-filtered (200 μM filter, Whatman/GE Healthcare). Afterwards, the herbal infusions were stored in aliquots at −80°C. For the infusions based on dried herbs, 3 g of material per 100 ml were utilized. The 10-min infusion was prepared by boiling up dried sage leaves in water (30 g per liter) and subsequent incubation for 10 min before the herb was removed. Sterile-filtered aliquots were then stored at −80°C. Infusions of frozen and re-thawed herbal material were prepared in the same manner as those based on fresh herbs. The concentration of 150 g of material per liter was calculated based on the fresh weight before freezing. The sources of supply for the herbal leaves and plants are as follows: coriander and sage, farmer’s market (Essen, Germany); red and green perilla plants, online vendor Naturkraeutergarten (Kleinich, Germany); bi-color perilla plant, home-grown; dried red perilla, home-dried; dried green perilla, Keiko Shiso Finest Selection (Japan); dried sage and dried thyme, vom-Achterhof Bio-Salbei/Bio-Thymian (Uplengen, Germany).

### in-cell-ELISA (icELISA)

For the quantification of viral protein amounts in infected cells, an icELISA was applied. A detailed icELISA protocol is provided in ^29^. Briefly, cells were infected with SARS-CoV-2 and fixed after 20 or 30 h of infection with 4% (w/v) paraformaldehyde/PBS. Cells were permeabilized with 1% (v/v) Triton-X-100/PBS and blocked with 3% (v/v) FCS/PBS. The primary antibody was added and incubated for 2 h at room temperature (RT) or over-night at 4°C. Peroxidase-labelled secondary antibody was incubated for 1-2 h. Washing steps were performed with 0.05% (v/v) Tween-20/PBS. Tetramethylbenzidin (TMB) substrate was added to visualize the enzyme reaction. The reaction was stopped with 0.5 M HCl before the absorbance was determined using a microplate multireader and MicroWin software (Mithras2 LB 943; Berthold Technologies). The resulting data were analyzed using Excel and GraphPad Prism software. The α-S mAb (kindly provided by Peter Miethe, fzmb, Bad Langensalza, Germany), α-N mAb (ABIN6952435), and POD-coupled secondary antibodies (Dianova) were used.

### Dose-response curves of antiviral activity

To enable comparison among different icELISA measurements and experiments, we included on every plate a virus calibration curve. Residual infectivity after treatment was calculated using the formula computed from the calibration curve (see Supplementary Fig. 1 as an example). Dose-response curves were compiled based on the relative change in infectivity compared to the untreated control.

### Immunofluorescence microscopy

Cells were infected with SARS-CoV-2 and fixed after 20 or 30 h of infection using 4% (w/v) paraformaldehyde/PBS for >2 h before they were discharged from the BSL-3 laboratory. Cells were permeabilized with 1% (v/v) Triton-X-100/PBS and blocked with 3% (v/v) FCS/PBS. SARS-CoV-2 infection was visualized by use of α-S mAb (kindly provided by Peter Miethe, fzmb, Bad Langensalza, Germany) and Cy2-conjugated goat anti-mouse IgG (Dianova). Nuclei were counterstained with 4’6-diamidino-2-phenylindole (DAPI; Sigma). Fluorescence was visualized using a THUNDER Imager 3D Cell Culture (Leica). Image analysis and processing were performed with LAS X Premium imaging software (Leica).

### Quantitative reverse-transcription PCR (qRT-PCR)

SARS-CoV-2 progeny was analyzed by quantification of viral RNA extracted from the culture supernatants of infected cells using the INSTANT Virus RNA/DNA Kit (Analytik Jena, Germany). Viral RNA was quantified by diagnostic qRT-PCR targeting the SARS-CoV-2 genes S and E (RealStar ^®^ SARS-CoV-2 RT-PCR kit, Altona, Hamburg, Germany). In the case of Omicron, undiluted RNA preparations were used as template for qRT-PCR, for all other variants, a 1:50 dilution was used.

### Interferons, inhibitors, substances, and size exclusion

Human IFNα2 was purchased from PBL Assay Science (#11101) and IFNβ from Peprotech (#300-02BC). Remdesivir and ruxolitinib were obtained from Cayman Chemicals (#30354) and Cell Guidance Systems (#SM87-10), respectively. The substances cinnamic acid, hydroxy-cinnamic acid, and dihydroxy-cinammic acid/caffeic acid were purchased from Sigma (#8002350250, #8002370050, #8220290010). Perilla aldehyde and perillyl alcohol were also obtained from Sigma (#W355704 and #218391). Size exclusion and protein fractionation of components of herbal infusions were conducted by use of Amicon 100K, 30K, 10K, and 3K filters (Sigma #UFC510024, #UFC501096, #UFC503096, #Z740183-96EA). The fraction of proteins >1 kDa was obtained by dialysis using the Mini Dialysis Kit 1 kDa (Sigma #GE80-6483-94). All kits were applied according to the manufacturer’s instructions.

### Sample preparation for MS

For protein extract preparation, 80-90 % confluent cell monolayers of a T25 flask of Vero E6 or Caco-2 cells were harvested using a cell scraper. Medium and cells were transferred to a 15 ml tube and centrifuged for 3 minutes at 350 g. Pellets were washed twice with 10 ml PBS, resuspended in 400 μl of lysis buffer (50 mM Tris, 150 mM NaCl, 1% [w/v] SDS, pH 7.8 supplemented with one cOmplete mini EDTA-free protease inhibitor tablet as well as one PhosSTOP tablet [both Roche] per 10 ml volume of lysis buffer), and transferred into 1.5 ml tubes. Lysates were incubated for 30 min at 4°C before they were subjected to virus inactivation at 70°C for 10 min followed by 10 min at 95°C. After ultrasonic treatment for 1 min, cell lysates were stored at −80°C until further analysis. 6 μl benzonase (27 units/μl, Merck) and 1 μl MgCl_2_ (2.5 mM final concentration) were added per sample, followed by incubation at 37°C for 30 min. Protein concentration was determined by a bicinchoninic acid assay (Pierce) and 100 μg protein lysate per replicate (n = 5 for all analyzed conditions) was processed further as described below.

### Positive pressure filter-aided sample preparation (FASP) in 96-well format

Cysteines were reduced using 10 mM dithiothreitol (Roche) at 56°C for 30 min and alkylated in the presence of 25 mM iodoacetamide (Sigma) for 30 min at RT in the dark. Samples were diluted at least 1:4 in 8 M urea (Sigma-Aldrich) dissolved in 100 mM Tris-HCl, pH 8.5 (Applichem) and transferred to a 30 kDa AcroPrep Omega filter membrane plate (PALL, New York, US, purchased via VWR, Hannover, Germany, REF 8035/518-0028) in a blocked randomized order, which was also kept for subsequent LC-MS measurements. The filter plate was placed on top of a 2.2 ml MegaBlock collection plate (Sarstedt, Nümbrecht, Germany) and the liquid of the protein solution was forced through the filter using a Resolvex A200 (Tecan, Männedorf, Switzerland) connected to nitrogen gas (N_2_, 5.5 bar, purity 4.8 or higher, Linde, Düsseldorf, Germany) using a relative pressure of 20% of the low profile setting. Subsequently, the dispensing function of the A200 was used to wash the filter twice with 200 μl 8 M urea buffer in Tris-HCl pH 8.5 and twice with 200 μl 50 mM ammonium bicarbonate (ABC, Fluka). After each washing step, the liquid was forced through the filter using the same pressure profile as for loading. Afterwards, the plate was centrifuged for 2 min at RT and 1,000 g to remove residual drops under the membrane. For digestion, 100 μl digestion buffer were added comprising 100 mM urea, 50 mM ABC, 2 mM CaCl_2_ (Merck) including sequencing grade trypsin (Promega, sequencing grade modified trypsin) in a concentration to meet a 1:3 (w/w) enzyme-to-sample ratio. After incubation for 16 h at 37°C, the digested protein fraction was forced through the filter and collected in a 500 μl LoBind plate (Eppendorf, Hamburg, Germany). The filter was further washed with 100 μl 50 mM ABC followed by 100 μl H_2_O, both wash steps were also collected in the LoBind plate. Then, tryptic digestion was stopped by reducing the pH <2 through the addition of trifluoroacetic acid (TFA, Biosolve, Valkenswaard, Netherlands) to a final concentration of 1% (v/v). Aliquots were transferred to 700 μl glass-vial plates (Waters, Eschborn, Germany) for injection on a monolithic column HPLC (for tryptic digestion quality control) ^73^. Based on the results, 4-5 replicates per condition were further analyzed by LC-MS.

### LC-MS in data-dependent acquisition-mode (DDA)

LC-MS was conducted using an UltiMate 3000 RSLCnano ProFlow UPLC system operated by Chromeleon Client 6.80 and online-coupled to a Q Exactive HF MS operated by Tune application 2.8 SP1 and Thermo Scientific Xcalibur 3.0.63 (both Thermo Scientific, Dreieich, Germany, including HPLC columns). Employed solvents were LC-MS grade or higher (Biosolve, Valkenswaard, Netherlands). 825 ng of tryptic peptides per LC-MS injection were analyzed. Samples were loaded on a trapping column (Acclaim PepMap C18, 0.1 x 20 mm, 5 μm, 100 Å) for 3 min in 0.1% TFA at a flow rate of 30 μl/min. Then, the trapping column was switched in line with the analytical column (Acclaim PepMap C18; 0.075 x 500 mm, 2 μm, 100 Å) and peptides were separated at a flow rate of 250 nl/min using a 102 min linear gradient of buffer B (84% v/v acetonitrile [ACN], 0.1% v/v formic acid [FA]) in buffer A (0.1% v/v FA) ranging from 3 to 25% B, followed by a 10 min linear gradient ranging from 25 to 35% B, washing steps and reconditioning of the analytical column to 3% B. Both columns were kept at 60°C temperature.

The UPLC system was coupled to the Q Exactive HF MS via a NSI Source (Thermo). Coated emitters (Silica Tip, 20 μm inner diameter, 10 μm tip inner diameter, New Objectives, Woburn, MA) and a static voltage of 1.8 kV were applied for electrospray ionization, ion transfer tube temperature was set to 250°C. The MS was operated in data-dependent acquisition (DDA) mode at positive polarity; all spectra were acquired in profile mode with survey scans acquired at a resolution of 60,000 followed by 15 MS/MS scans at a resolution of 15,000 (top15). Precursor ions were selected for MS/MS by intensity, isolated in a 1.6 m/z window and subjected to fragmentation by higher energy collision induced dissociation using a normalized collision energy (NCE) of 27. Automatic gain control target values were set to 10^6^ and 5×10^4^ and the maximum ion injection was set to 120 ms and 50 ms for MS and MS/MS, respectively. Precursor masses were excluded from re-fragmentation for 20 sec (dynamic exclusion).

### Protein identification and relative quantification with Proteome Discoverer

DDA files were processed with Proteome Discoverer 2.4 (Thermo Scientific) using Spectrum Files RC and Sequest HT nodes as database search algorithm and Percolator ^74^ in conjunction with Peptide validator and Protein FDR validator nodes for adjusting the false discovery rate to 1% on PSM, peptide, and protein levels. Database search was conducted against Uniprot homo sapiens database for Caco-2 (UP000005640, 20,376 entries, retrieved in November 2019) or Chlorocebus sabaeus database for Vero E6 (UP000029965, 19,230 entries, retrieved in January 2021) in conjunction with the SARS-CoV-2 database (UP000464024, 14 entries, retrieved in January 2021) supplemented with reported putative novel ORFs ^75^, common contaminants and an additional fasta file containing the amino acid sequence of indexed retention time peptides added as an internal standard ^76^ by the Sequest HT search engine with the following parameters: error tolerances of 10 ppm and 0.02 Da for precursor and fragment, trypsin (full) as enzyme with a maximum of two missed cleavage sites, oxidation of Met as variable modification (+15.995 Da) and carbamidomethylation of Cys (+57.021 Da) as fixed modification. Full settings are provided as supplementary material via ProteomeXchange. Quantification was performed using the Minora feature detector node in conjunction with the Feature mapper node with a maximum retention time shift of 10 min and mass tolerance of 10 ppm for chromatographic alignment and minimum signal to noise threshold of 10 for feature linking and mapping. Precursor Ions Quantifier node accepted only unique peptides for quantification while considering protein groups for peptide uniqueness with disabled scaling and normalization to total peptide amount. Sample abundances of the connected peptide groups were summed to calculate protein abundances. Common contaminant proteins and proteins with less than two unique peptides per protein group were filtered out. Normalized protein abundances were used for further data analysis.

### Analysis of MS data

For all analyses, only proteins with a maximum of one missing replicate value per respective condition were considered. Lightweight normalizing and testing tool (LNTT) ^77^ was used for the creation of volcano plots in Fig. 5 and Supplementary Fig. 10. Proteome Discoverer normalized (total protein normalization) data were filtered (at least two unique peptides per protein group required) and proteins with a coefficient of variation <20.0% over all measurements (also across treatments) were removed. Independent two-sided two-sample Welch’s t-test was performed for remaining proteins. Computed p values were adjusted using the Benjamini and Hochberg multiple test correction (https://www.jstor.org/stable/2346101) and results were plotted by LNTT. Gene names for labeling were retrieved by accession numbers using Uniprot API. For heatmap analysis, only proteins quantified across all conditions and time points were considered for Vero E6 and Caco-2 samples, respectively. Displayed order of the proteins was sorted accordingly to the mean value for the respective 6 h p.i.

### Statistical analysis

Statistical significance was determined using one-way ANOVA as described in the figure legends. A p value of <0.05 was considered statistically significant. *, p value <0.05. **, p value <0.01. ***, p value <0.001. Calculation of p values for analysis of MS data are described above.

### Data availability statement

Data generated or analyzed during this study are included in this published article and its supplementary information files. The MS proteomics data have been deposited to the ProteomeXchange Consortium ^78^ via the PRIDE ^79^ partner repository with the dataset identifier PXD030742 (user, password: vjMdjgnz).

## Acknowledgements

We thank Cornelia Schumbrutzki for excellent technical support and the parents of VTKLT for providing the bi-color perilla plants. ISAS acknowledges the support by the Ministerium für Kultur und Wissenschaft des Landes Nordrhein-Westfalen, the Regierender Bürgermeister von Berlin - inkl. Wissenschaft und Forschung, and the Bundesministerium für Bildung und Forschung (313-8.04.10.02-149281). AS receives funding from the Deutsche Forschungsgemeinschaft (DFG) through TRR 240 Z02. The members of the Institute for Virology Essen are very grateful for a very generous donation from Alantra, based on which the Leica THUNDER Imager was purchased.

## Author contributions

VTKLT conceived the project, designed and performed experiments, analyzed data, prepared figures, and wrote the manuscript. DM, CS, YFM, BK, KW performed experiments. RS, JKL conducted MS analyses. RS, JKL, DK analyzed MS data. SL developed methodology for sample preparation. FM analyzed NGS data. MG, UD, AS designed experiments. MT conceived the project, designed and performed experiments, analyzed data, and wrote the manuscript. All authors reviewed and edited the manuscript.

## Supplementary information

Supplementary Fig. 1: Representative example of the virus calibration curves (included on every plate) applied to calculate the residual infectivity after treatment.

**a** icELISA data (α-S staining) of SARS-CoV-2-infected Vero E6 cells after treatment with herbal infusions. Each condition was analyzed in triplicate.

**b** icELISA data of the virus calibration curve. Each condition was analyzed in duplicate.

**c** The formula of the calibration curve from the same plate was applied to calculate the residual PFU after treatment.

Supplementary Fig. 2: Two different clinical SARS-CoV-2 isolates exhibit almost identical susceptibilities towards perilla and sage.

**a, b** Pooled icELISA data of 3 (**a**, SARS-CoV-2 variant B.1) and 5 (**b**, SARS-CoV-2 variant B.1.1.232) independent experiments of SARS-CoV-2-infected Vero E6 cells after treatment with herbal infusions and icELISA using α-S or α-N mAbs for staining. Data are expressed as relative change in optical density compared to the untreated control. The perilla- and sage-treated conditions were compared to the corresponding coriander-treated condition (same dilution) by one-way ANOVA. **, p<0.01. ***, p<0.001.

Supplementary Fig. 3: Preserved sage and perilla leaves retain bioactive compounds.

**a** The one-hour treatment with the herbal infusions is not cytotoxic. Vero E6 cells were treated with indicated dilutions of herbal infusions in parallel to an infection experiment. At 18 h post treatment, cell viability was analyzed by Orangu cell counting solution (Cell Guidance Systems).

**b** Representative dose-response curves of SARS-CoV-2-infected Vero E6 cells (2000 PFU per well) after treatment with herbal infusions. Upper panel depicts the results for 1 h of treatment, lower panel the results for 30 min of treatment. Data are expressed as relative change in infectivity compared to the untreated control. Each condition was analyzed in triplicate. The perilla- and sage-treated conditions were compared to the corresponding coriander-treated condition (same dilution) by one-way ANOVA. **, p<0.01. ***, p<0.001.

**c** Representative dose-response curves of SARS-CoV-2-infected Vero E6 cells after treatment with aqueous infusions generated from fresh or dried red perilla, green perilla, or sage for 1 h. Each condition was analyzed in triplicate. All conditions were compared to the untreated control by one-way ANOVA. *, p<0.05. **, p<0.01. ***, p<0.001.

**d** Representative dose-response curves of SARS-CoV-2-infected Vero E6 cells after treatment with herbal infusions generated by over-night or 10-min extraction of dried sage (see methods section for details). Each condition was analyzed in triplicate. All conditions were compared to the untreated control by one-way ANOVA. ***, p<0.001.

**e** Representative dose-response curves of SARS-CoV-2-infected Vero E6 cells after treatment with aliquots of herbal infusions that were thawed once or twice. Each condition was analyzed in triplicate. All conditions were compared to the untreated control by one-way ANOVA. *, p<0.05. ***, p<0.001.

**f** Representative dose-response curves of SARS-CoV-2-infected Vero E6 cells after treatment with aqueous infusions generated from fresh or frozen red or green perilla. Each condition was analyzed in triplicate. All conditions were compared to the untreated control by one-way ANOVA. ***, p<0.001.

Supplementary Fig. 4: Perilla and sage elicit prophylactic antiviral activity *in vitro.*

**a** Scheme of the experimental setup for the *in vitro* analysis of prophylactic effects against SARS-CoV-2.

**b** Pooled icELISA data of 6 independent experiments of SARS-CoV-2-infected Vero E6 cells after prophylactic treatment with herbal infusions using two distinct SARS-CoV-2 isolates (B.1 and B. 1.1.232) for infection of Vero E6 and α-S or α-N mAbs for staining. Data are expressed as relative change in optical density compared to the untreated control. The perilla- and sage-treated conditions were compared to the corresponding coriander-treated condition (same dilution) by one-way ANOVA. *, p<0.05. **, p<0.01. ***, p<0.001.

**c, d** Comparison of prophylactic and therapeutic treatment (−1 to 0 h p.i. verus 1 to 2 h p.i.) of SARS-CoV-2-infected Vero E6 cells (2000 PFU per well). Each condition was analyzed in triplicate. All conditions were compared to the untreated control by one-way ANOVA. **, p<0.01. ***, p<0.001.

Supplementary Fig. 5: Perilla and sage confer protection against SARS-CoV-2 infection in human cells.

**a** Scheme of the applied experimental setup.

**b** icELISA data (α-S staining) of SARS-CoV-2-infected Caco-2 cells (2000 PFU per well). Data are expressed as relative change in optical density compared to the untreated control. Each condition was analyzed in triplicate.

Supplementary Fig. 6: Size exclusion and protein fractionation analyses of perilla infusion disclosed compounds <10 kDa as active components.

Pooled icELISA data (α-S staining) of 2 independent experiments of SARS-CoV-2-infected Vero E6 cells after treatment with indicated fractions of perilla infusion. Data are expressed as relative change in optical density compared to the untreated control. Protein fractionation of perilla infusion components was conducted by use of Amicon 100K, 30K, and 10K filters. Each condition was analyzed in triplicate. Dilutions and concentrations by the fractionation steps were considered. All conditions were compared to the untreated control by one-way ANOVA. ***, p<0.001.

**a** Perilla flow-through of Amicon 100K, 30K, and 10K filters were analyzed.

**b** Perilla fractions >100 kDa, >30 kDa, and >10 kDa were analyzed.

Supplementary Fig. 7: Dihydroxy-cinnamic acid, perilla aldehyde, and perillyl alcohol are antiviral components of herbal infusions.

**a, b** icELISA data of SARS-CoV-2-infected Caco-2 cells after treatment with indicated substances at 1.5 h p.i. for 1.5 h. Data are expressed as relative change in optical density compared to the untreated control. Each condition was analyzed in triplicate. All treated conditions were compared to the untreated control by one-way ANOVA. *, p<0.05. ***, p<0.001.

**c** Pooled icELISA data of SARS-CoV-2-infected cells (Vero E6, 4 independent experiments; Caco-2, 3 independent experiments) after treatment with perilla or thyme infusion. Data are expressed as relative change in optical density compared to the untreated control. All treated conditions were compared to the untreated control by one-way ANOVA. *, p<0.05. **, p<0.01. ***, p<0.001.

Supplementary Fig. 8: Herbal infusion-induced changes of proposed host restriction and host dependency factors in Vero E6 cells.

SARS-CoV-2-infected Vero E6 cells as well as mock controls were treated at 1 h p.i. with perilla and sage infusion for 1 h. At 2, 6, and 20 h p.i., cells were lysed and subjected to global MS analysis. Each condition was analyzed in quintuplicate. Proposed HRFs and HDFs ^33-35^, which were consistently quantified across all investigated conditions, were analyzed regarding the changes induced by treatment with perilla or sage infusions. The ratio of the treated condition to the respective untreated condition is shown as log_2_ fold change.

Supplementary Fig. 9: Herbal infusion-induced changes of proposed host restriction and host dependency factors in Caco-2 cells.

SARS-CoV-2-infected Caco-2 cells as well as mock controls were treated at 1 h p.i. with perilla and sage infusion for 1 h. At 2, 6, and 30 h p.i., cells were lysed and subjected to global MS analysis. Each condition was analyzed in quadruplicate or quintuplicate. Proposed HRFs and HDFs ^33-35^, which were consistently quantified across all investigated conditions, were analyzed regarding the changes induced by treatment with perilla or sage infusions. The ratio of the treated condition to the respective untreated condition is shown as log_2_ fold change.

Supplementary Fig. 10: Perilla and sage infusion-induced changes at 6 h p.i. in infected Caco-2 and Vero E6 cells.

**a, b** Volcano plots of MS results obtained from infected Caco-2 (**a, b**) or Vero E6 (**c, d**) at 6 h p.i. Proteome Discoverer normalized data were filtered (at least two unique peptides per protein group required) and proteins with a coefficient of variation <20.0% over all measurements (also across treatments) were removed (see methods section for details).

Supplementary Fig. 11: Herbal infusions exhibit antiviral activity against SARS-CoV-2 variants of concern.

**a** Vero E6 cells were infected with graded doses of SARS-CoV-2 wt and Beta (B.1.351). At 20 h p.i., cells were fixed and analyzed by icELISA using different antibodies recognizing the receptor-binding domain of Spike (RBD) or the nucleocapsid protein (N).

**b** Representative dose-response curves of Vero E6 cells infected with SARS-CoV-2 wt and variants of concern (Alpha/B.1.1.7 and Beta/B.1.351) after treatment with aqueous infusions of coriander, perilla, or sage at 1 h p.i. for 1 h. SARS-CoV-2 replication was analyzed at 20 h p.i. by icELISA (α-N staining). Data are expressed as relative change in infectivity compared to the untreated control. Each condition was analyzed in triplicate. The comparison of the herb-treated samples of SARS-CoV-2 to the untreated control by one-way ANOVA showed for all dilutions of coriander no significance. The perilla- and sage-treated conditions of SARS-CoV-2 were also compared to the corresponding coriander-treated condition (same dilution) and these results are depicted in the diagram. *, p<0.05. **, p<0.01. ***, p<0.001.

Supplementary File 1: Microsoft Excel-based plotting option for relative abundances of all individual proteins detected in Vero E6 samples by at least two unique peptides

Supplementary File 2: Microsoft Excel-based plotting option for relative abundances of all individual proteins detected in Caco-2 samples by at least two unique peptides

## References

1. Solecki, R.S. Shanidar IV, a Neanderthal Flower Burial in Northern Iraq. Science 190, 880–881 (1975).

2. Lietava, J. Medicinal plants in a Middle Paleolithic grave Shanidar IV? J Ethnopharmacol 35, 263–266 (1992).

3. Lu, H., et al. Earliest tea as evidence for one branch of the Silk Road across the Tibetan Plateau. Sci Rep 6, 18955 (2016).

4. Yu, H., et al. Phytochemical and phytopharmacological review of Perilla frutescens L. (Labiatae), a traditional edible-medicinal herb in China. Food Chem Toxicol 108, 375–391 (2017).

5. Beheshti-Rouy, M., et al. The antibacterial effect of sage extract (Salvia officinalis) mouthwash against Streptococcus mutans in dental plaque: a randomized clinical trial. Iran J Microbiol 7, 173–177 (2015).

6. Kim, D.H., Kim, Y.C. & Choi, U.K. Optimization of antibacterial activity of Perilla frutescens var. acuta leaf against Staphylococcus aureus using evolutionary operation factorial design technique. Int J Mol Sci 12, 2395–2407 (2011).

7. Yuan, H., Ma, Q., Ye, L. & Piao, G. The Traditional Medicine and Modern Medicine from Natural Products. Molecules 21(2016).

8. Tu, Y. Artemisinin-A Gift from Traditional Chinese Medicine to the World (Nobel Lecture). Angew Chem Int Ed Engl 55, 10210–10226 (2016).

9. Krishna, S., Bustamante, L., Haynes, R.K. & Staines, H.M. Artemisinins: their growing importance in medicine. Trends Pharmacol Sci 29, 520–527 (2008).

10. Zhu, N., et al. A Novel Coronavirus from Patients with Pneumonia in China, 2019. N Engl J Med 382, 727–733 (2020).

11. Coronaviridae Study Group of the International Committee on Taxonomy of, V. The species Severe acute respiratory syndrome-related coronavirus: classifying 2019-nCoV and naming it SARS-CoV-2. Nat Microbiol 5, 536–544 (2020).

12. Wang, S., et al. A Crowned Killer’s Resume: Genome, Structure, Receptors, and Origin of SARS-CoV-2. Virol Sin (2020).

13. Liu, J., et al. Overlapping and discrete aspects of the pathology and pathogenesis of the emerging human pathogenic coronaviruses SARS-CoV, MERS-CoV, and 2019-nCoV. J Med Virol 92, 491–494 (2020).

14. Wiersinga, W.J., Rhodes, A., Cheng, A.C., Peacock, S.J. & Prescott, H.C. Pathophysiology, Transmission, Diagnosis, and Treatment of Coronavirus Disease 2019 (COVID-19): A Review. JAMA 324, 782–793 (2020).

15. Gupta, A., et al. Extrapulmonary manifestations of COVID-19. Nat Med 26, 1017–1032 (2020).

16. Yuki, K., Fujiogi, M. & Koutsogiannaki, S. COVID-19 pathophysiology: A review. Clin Immunol 215, 108427 (2020).

17. Bar-On, Y.M., Flamholz, A., Phillips, R. & Milo, R. SARS-CoV-2 (COVID-19) by the numbers. Elife 9(2020).

18. Li, J., Lai, S., Gao, G.F. & Shi, W. The emergence, genomic diversity and global spread of SARS-CoV-2. Nature 600, 408–418 (2021).

19. Dong, E., Du, H. & Gardner, L. An interactive web-based dashboard to track COVID-19 in real time. Lancet Infect Dis 20, 533–534 (2020).

20. Nicola, M., et al. The socio-economic implications of the coronavirus pandemic (COVID-19): A review. Int J Surg 78, 185–193 (2020).

21. Menachery, V.D., et al. A SARS-like cluster of circulating bat coronaviruses shows potential for human emergence. Nat Med 21, 1508–1513 (2015).

22. Lau, S.K., et al. Severe acute respiratory syndrome coronavirus-like virus in Chinese horseshoe bats. Proc Natl Acad Sci U S A 102, 14040–14045 (2005).

23. Li, W., et al. Bats are natural reservoirs of SARS-like coronaviruses. Science 310, 676–679 (2005).

24. Lednicky, J.A., et al. Independent infections of porcine deltacoronavirus among Haitian children. Nature 600, 133–137 (2021).

25. Vijgen, L., et al. Complete genomic sequence of human coronavirus OC43: molecular clock analysis suggests a relatively recent zoonotic coronavirus transmission event. J Virol 79, 1595–1604 (2005).

26. Moreira-Soto, A., et al. Neotropical Bats from Costa Rica harbour Diverse Coronaviruses. Zoonoses Public Health 62, 501–505 (2015).

27. Ahmed, I., et al. Behavioral preventive measures and the use of medicines and herbal products among the public in response to Covid-19 in Bangladesh: A cross-sectional study. medRxiv, 2020.2008.2015.20175513 (2020).

28. Alyami, H.S., et al. Knowledge about COVID-19 and patients beliefs about and use of herbal products during the COVID-19 pandemic: a cross-sectional study in Saudi Arabia. medRxiv, 2020.2006.2023.20138107 (2020).

29. Scholer, L., et al. A Novel In-Cell ELISA Assay Allows Rapid and Automated Quantification of SARS-CoV-2 to Analyze Neutralizing Antibodies and Antiviral Compounds. Front Immunol 11, 573526 (2020).

30. Hoffmann, M., et al. SARS-CoV-2 Cell Entry Depends on ACE2 and TMPRSS2 and Is Blocked by a Clinically Proven Protease Inhibitor. Cell 181, 271–280 e278 (2020).

31. Duelund, L., Amiot, A., Fillon, A. & Mouritsen, O.G. Influence of the active compounds of Perilla frutescens leaves on lipid membranes. J Nat Prod 75, 160–166 (2012).

32. Peng, Y., Ye, J. & Kong, J. Determination of phenolic compounds in Perilla frutescens L. by capillary electrophoresis with electrochemical detection. J Agric Food Chem 53, 8141–8147 (2005).

33. Wei, J., et al. Genome-wide CRISPR screen reveals host genes that regulate SARS-CoV-2 infection. bioRxiv (2020).

34. Daniloski, Z., et al. Identification of Required Host Factors for SARS-CoV-2 Infection in Human Cells. Cell 184, 92–105 e116 (2021).

35. Wang, R., et al. Genetic Screens Identify Host Factors for SARS-CoV-2 and Common Cold Coronaviruses. Cell 184, 106–119 e114 (2021).

36. Maines, M.D. Heme oxygenase: function, multiplicity, regulatory mechanisms, and clinical applications. FASEB J 2, 2557–2568 (1988).

37. Kikuchi, G., Yoshida, T. & Noguchi, M. Heme oxygenase and heme degradation. Biochem Biophys Res Commun 338, 558–567 (2005).

38. Connick, J.P., Reed, J.R., Cawley, G.F. & Backes, W.L. Heteromeric complex formation between human cytochrome P450 CYP1A1 and heme oxygenase-1. Biochem J 478, 377–388 (2021).

39. Jangra, S., et al. SARS-CoV-2 spike E484K mutation reduces antibody neutralisation. Lancet Microbe 2, e283–e284 (2021).

40. Barton, M.I., et al. Effects of common mutations in the SARS-CoV-2 Spike RBD and its ligand, the human ACE2 receptor on binding affinity and kinetics. Elife 10(2021).

41. Pan, H., et al. Repurposed antiviral drugs for COVID-19 –interim WHO SOLIDARITY trial results. medRxiv, 2020.2010.2015.20209817 (2020).

42. Hung, I.F., et al. Triple combination of interferon beta-1b, lopinavir-ritonavir, and ribavirin in the treatment of patients admitted to hospital with COVID-19: an open-label, randomised, phase 2 trial. Lancet 395, 1695–1704 (2020).

43. Diaz, M.O., et al. Homozygous deletion of the alpha- and beta 1-interferon genes in human leukemia and derived cell lines. Proceedings of the National Academy of Sciences 85, 5259–5263 (1988).

44. Newman, D.J. & Cragg, G.M. Natural Products as Sources of New Drugs over the Nearly Four Decades from 01/1981 to 09/2019. J Nat Prod 83, 770–803 (2020).

45. Khan, F.A., Maalik, A. & Murtaza, G. Inhibitory mechanism against oxidative stress of caffeic acid. J Food Drug Anal 24, 695–702 (2016).

46. Khan, F., Bamunuarachchi, N.I., Tabassum, N. & Kim, Y.M. Caffeic Acid and Its Derivatives: Antimicrobial Drugs toward Microbial Pathogens. J Agric Food Chem 69, 2979–3004 (2021).

47. Erhunmwunsee, F., et al. Recent development in biological activities and safety concerns of perillaldehyde from perilla plants: A review. Crit Rev Food Sci Nutr, 1–13 (2021).

48. Zhang, Y., et al. Perilaldehyde activates AMP-activated protein kinase to suppress the growth of gastric cancer via induction of autophagy. J Cell Biochem (2018).

49. Chen, T.C., Fonseca, C.O. & Schonthal, A.H. Preclinical development and clinical use of perillyl alcohol for chemoprevention and cancer therapy. Am J Cancer Res 5, 1580–1593 (2015).

50. Oh, C., et al. Inhibition of HIV-1 infection by aqueous extracts of Prunella vulgaris L. Virol J 8, 188 (2011).

51. Geuenich, S., et al. Aqueous extracts from peppermint, sage and lemon balm leaves display potent anti-HIV-1 activity by increasing the virion density. Retrovirology 5, 27 (2008).

52. Yamasaki, K., et al. Anti-HIV-1 activity of herbs in Labiatae. Biol Pharm Bull 21, 829–833 (1998).

53. Kawahata, T., et al. A novel substance purified from Perilla frutescens Britton inhibits an early stage of HIV-1 replication without blocking viral adsorption. Antivir Chem Chemother 13, 283–288 (2002).

54. Tay, M.Z., Poh, C.M., Renia, L., MacAry, P.A. & Ng, L.F.P. The trinity of COVID-19: immunity, inflammation and intervention. Nat Rev Immunol 20, 363–374 (2020).

55. Becker, R.C. COVID-19 update: Covid-19-associated coagulopathy. J Thromb Thrombolysis 50, 54–67 (2020).

56. Gibson, P.G., Qin, L. & Puah, S.H. COVID-19 acute respiratory distress syndrome (ARDS): clinical features and differences from typical pre-COVID-19 ARDS. Med J Aust 213, 54–56 e51 (2020).

57. Wichmann, D., et al. Autopsy Findings and Venous Thromboembolism in Patients With COVID-19: A Prospective Cohort Study. Ann Intern Med 173, 268–277 (2020).

58. Elezkurtaj, S., et al. Causes of death and comorbidities in hospitalized patients with COVID-19. Sci Rep 11, 4263 (2021).

59. Gong, Q., et al. Heme oxygenase-1 upregulation significantly inhibits TNF-alpha and Hmgb1 releasing and attenuates lipopolysaccharide-induced acute lung injury in mice. Int Immunopharmacol 8, 792–798 (2008).

60. Jin, Y., et al. Deletion of caveolin-1 protects against oxidative lung injury via up-regulation of heme oxygenase-1. Am J Respir Cell Mol Biol 39, 171–179 (2008).

61. Siner, J.M., et al. VEGF-induced heme oxygenase-1 confers cytoprotection from lethal hyperoxia in vivo. FASEB J 21, 1422–1432 (2007).

62. Maeshima, K., et al. Prevention of hemorrhagic shock-induced lung injury by heme arginate treatment in rats. Biochem Pharmacol 69, 1667–1680 (2005).

63. Otterbein, L.E., et al. Protective effects of heme oxygenase-1 in acute lung injury. Chest 116, 61S–63S (1999).

64. https://www.accessdata.fda.gov/scripts/cdrh/cfdocs/cfcfr/CFRSearch.cfm?fr=182.10.

65. https://www.cdc.gov/pregnancy/meds/treatingfortwo/facts.html.

66. Mayer, E., Gescheidt-Shoshany, H. & Weltfriend, S. Allergic contact dermatitis caused by Salvia officinalis extract. Contact Dermatitis 64, 237–238 (2011).

67. O’Toole, A., et al. Assignment of epidemiological lineages in an emerging pandemic using the pangolin tool. Virus Evol 7, veab064 (2021).

68. Rambaut, A., et al. A dynamic nomenclature proposal for SARS-CoV-2 lineages to assist genomic epidemiology. Nat Microbiol 5, 1403–1407 (2020).

69. Elbe, S. & Buckland-Merrett, G. Data, disease and diplomacy: GISAID’s innovative contribution to global health. Glob Chall 1, 33–46 (2017).

70. Shu, Y. & McCauley, J. GISAID: Global initiative on sharing all influenza data - from vision to reality. Euro Surveill 22(2017).

71. Julia L. Mullen, G.T., Alaa Abdel Latif, Manar Alkuzweny, Marco Cano, Emily Haag, Jerry Zhou, Mark Zeller, Emory Hufbauer, Nate Matteson, Kristian G. Andersen, Chunlei Wu, Andrew I. Su, Karthik Gangavarapu, Laura D. Hughes, and the Center for Viral Systems Biology. outbreak.info. Available online: https://outbreak.info/ (2020).

72. Farnsworth, A., Goldsmith, K. & Johnson, D.C. Herpes simplex virus glycoproteins gD and gE/gI serve essential but redundant functions during acquisition of the virion envelope in the cytoplasm. J Virol 77, 8481–8494 (2003).

73. Burkhart, J.M., Schumbrutzki, C., Wortelkamp, S., Sickmann, A. & Zahedi, R.P. Systematic and quantitative comparison of digest efficiency and specificity reveals the impact of trypsin quality on MS-based proteomics. J Proteomics 75, 1454–1462 (2012).

74. Kall, L., Canterbury, J.D., Weston, J., Noble, W.S. & MacCoss, M.J. Semi-supervised learning for peptide identification from shotgun proteomics datasets. Nat Methods 4, 923–925 (2007).

75. Finkel, Y., et al. The coding capacity of SARS-CoV-2. Nature 589, 125–130 (2021).

76. Escher, C., et al. Using iRT, a normalized retention time for more targeted measurement of peptides. Proteomics 12, 1111–1121 (2012).

77. Kopczynski, D., & Sakson, R. LNTT (Lightweight normalizing and testing tool) (1.0.1). Zenodo. https://doi.org/10.5281/zenodo.5566369 (2021).

78. Deutsch, E.W., et al. The ProteomeXchange consortium in 2020: enabling ‘big data’ approaches in proteomics. Nucleic Acids Res 48, D1145–D1152 (2020).

79. Perez-Riverol, Y., et al. The PRIDE database and related tools and resources in 2019: improving support for quantification data. Nucleic Acids Res 47, D442–D450 (2019).

