## Supplementary Fig. for "Universally available herbal teas based on sage and perilla elicit potent antiviral activity against SARS-CoV-2 variants of concern by HMOX-1 upregulation in human cells"

### Supplementary Figure 1

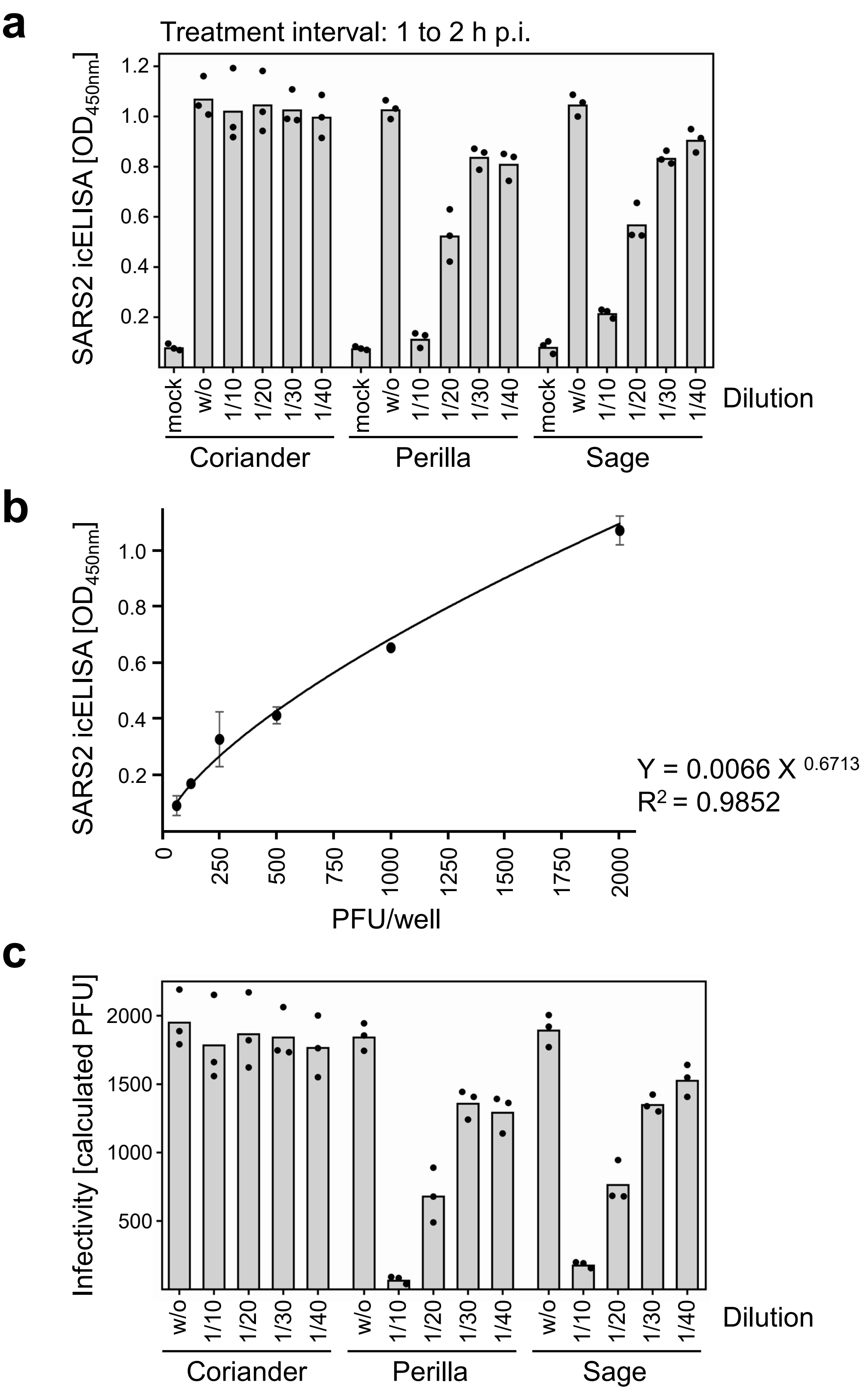

Supplementary Figure 2

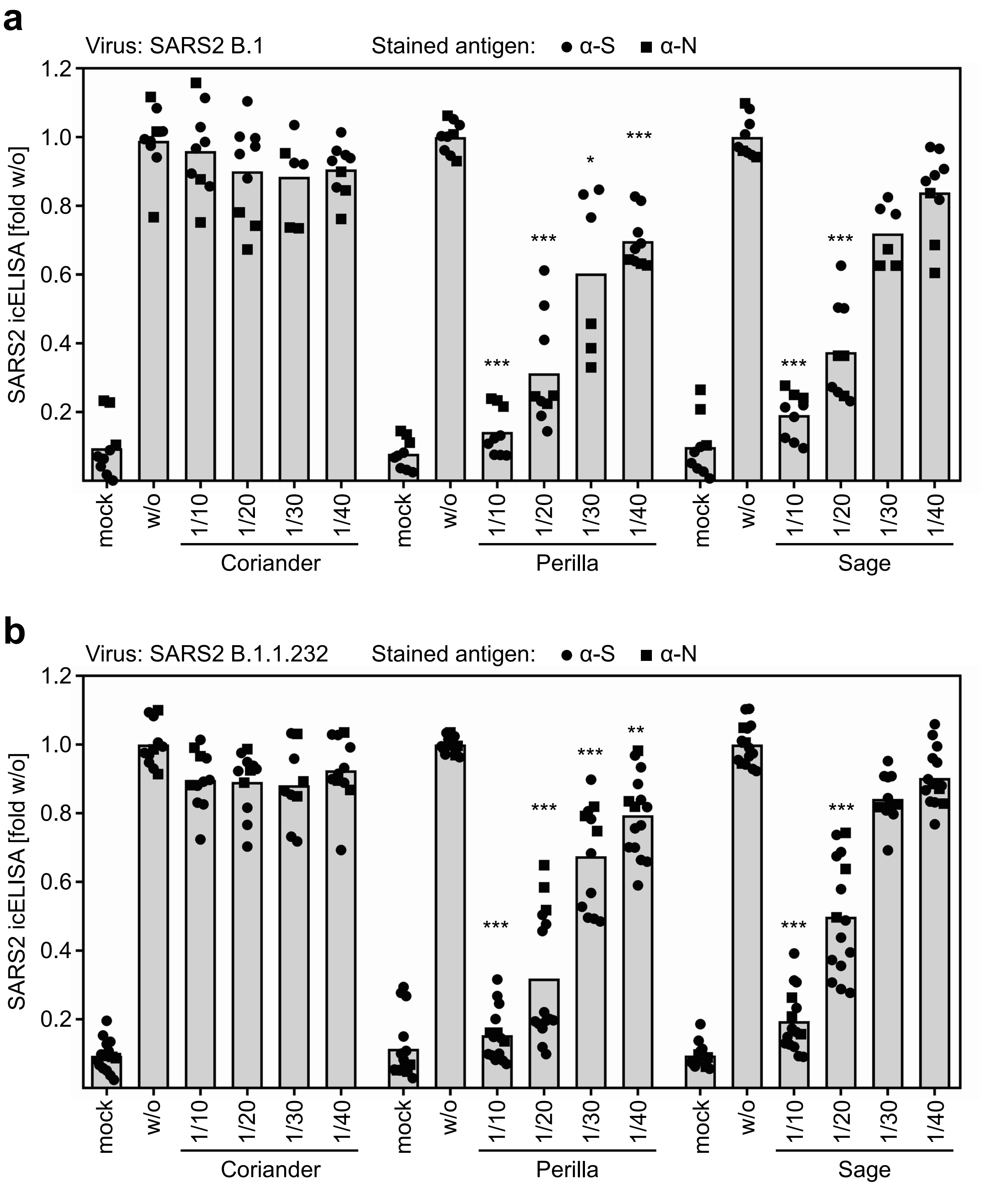

### Supplementary Figure 3

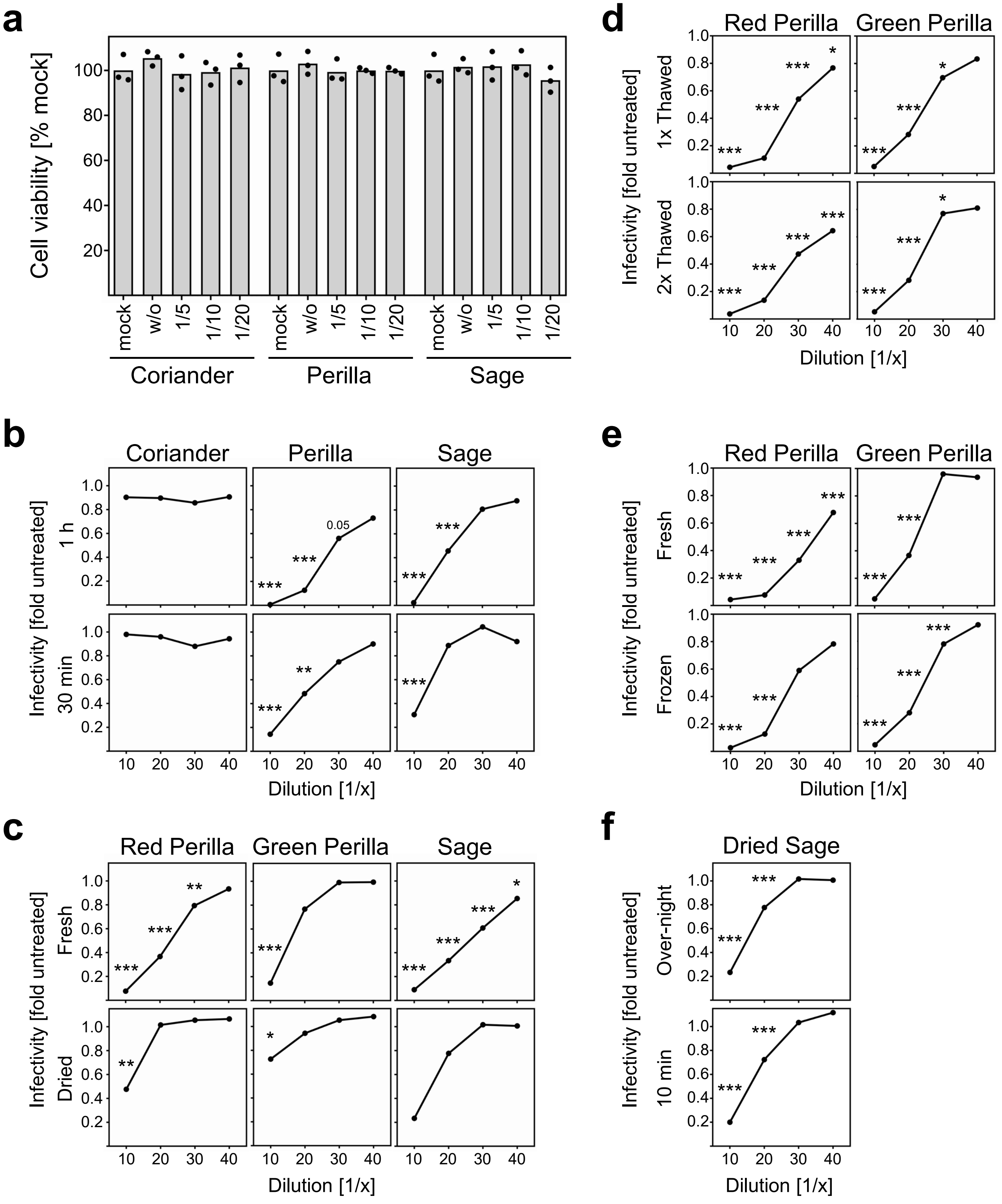

### Supplementary Figure 4

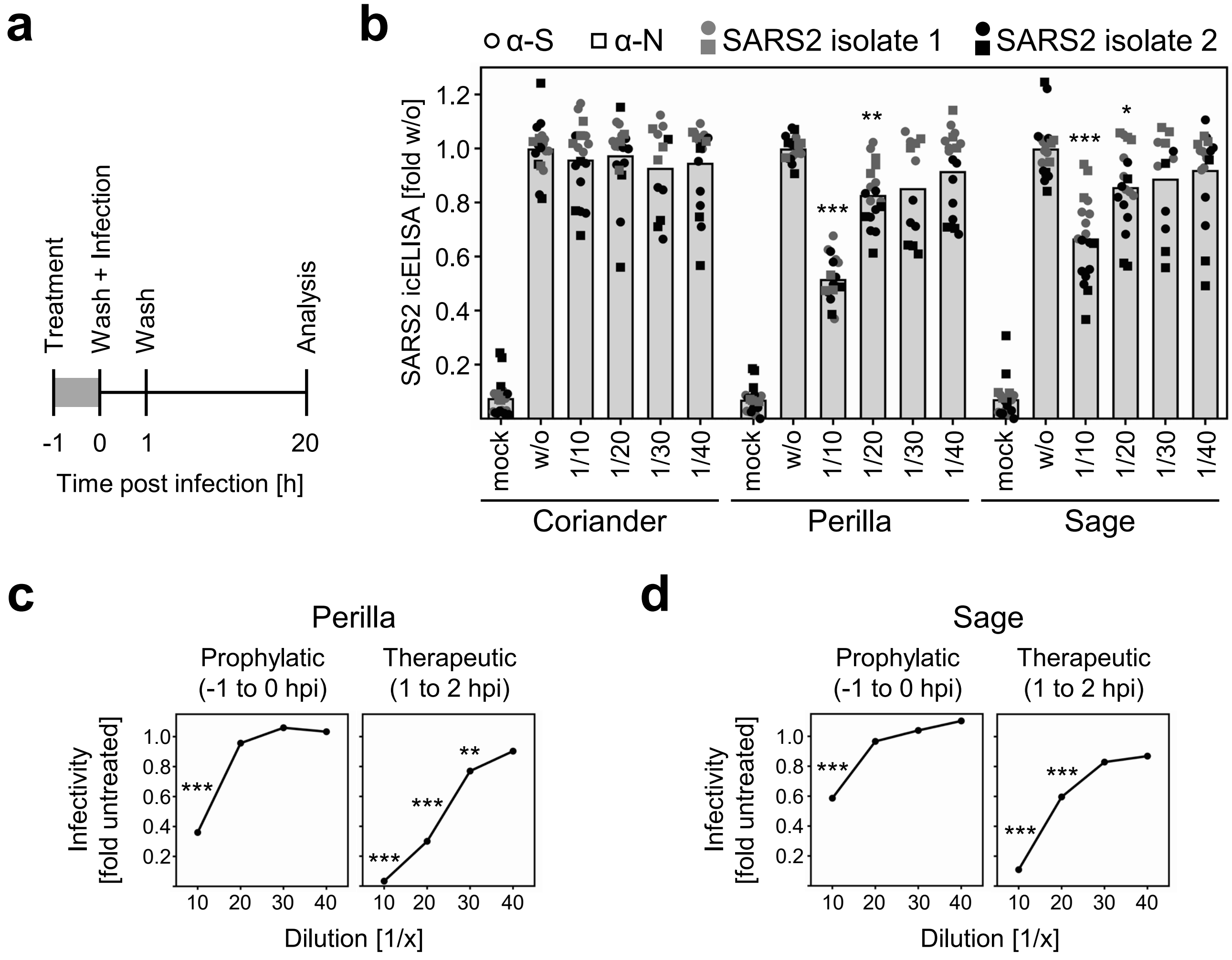

### Supplementary Figure 5

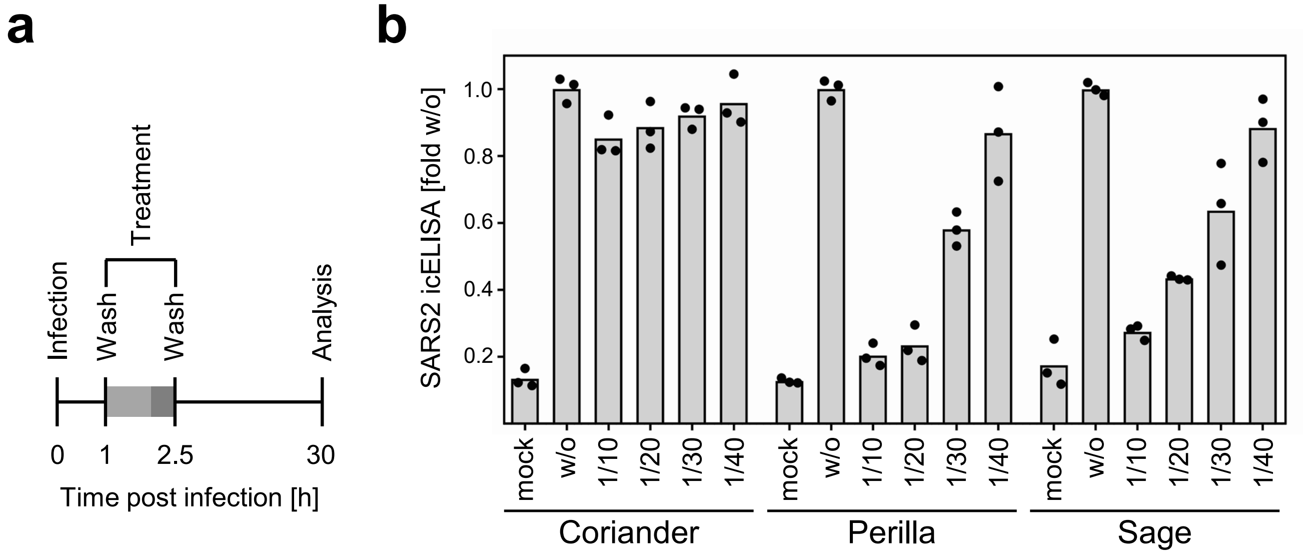

### Supplementary Figure 6

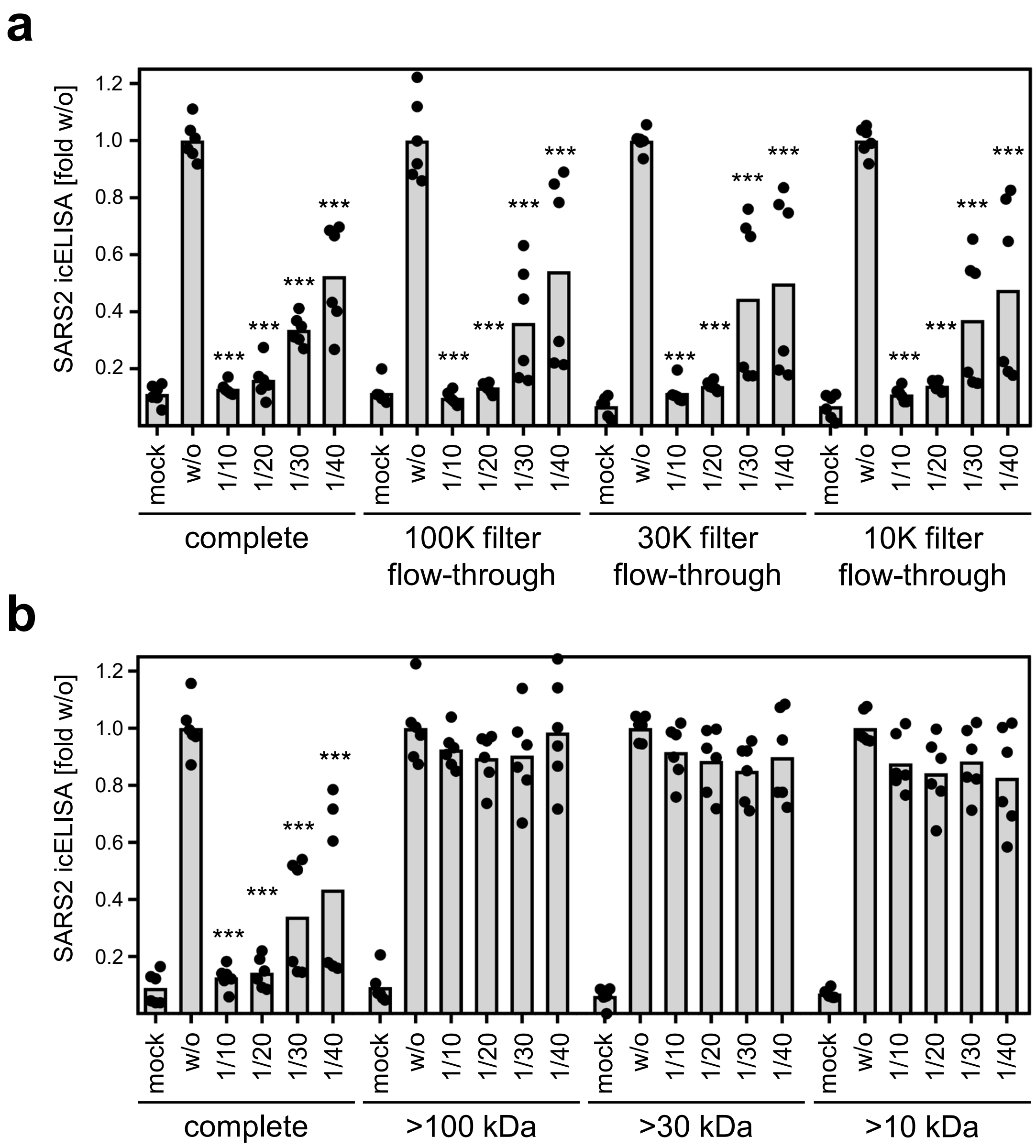

### Supplementary Figure 7

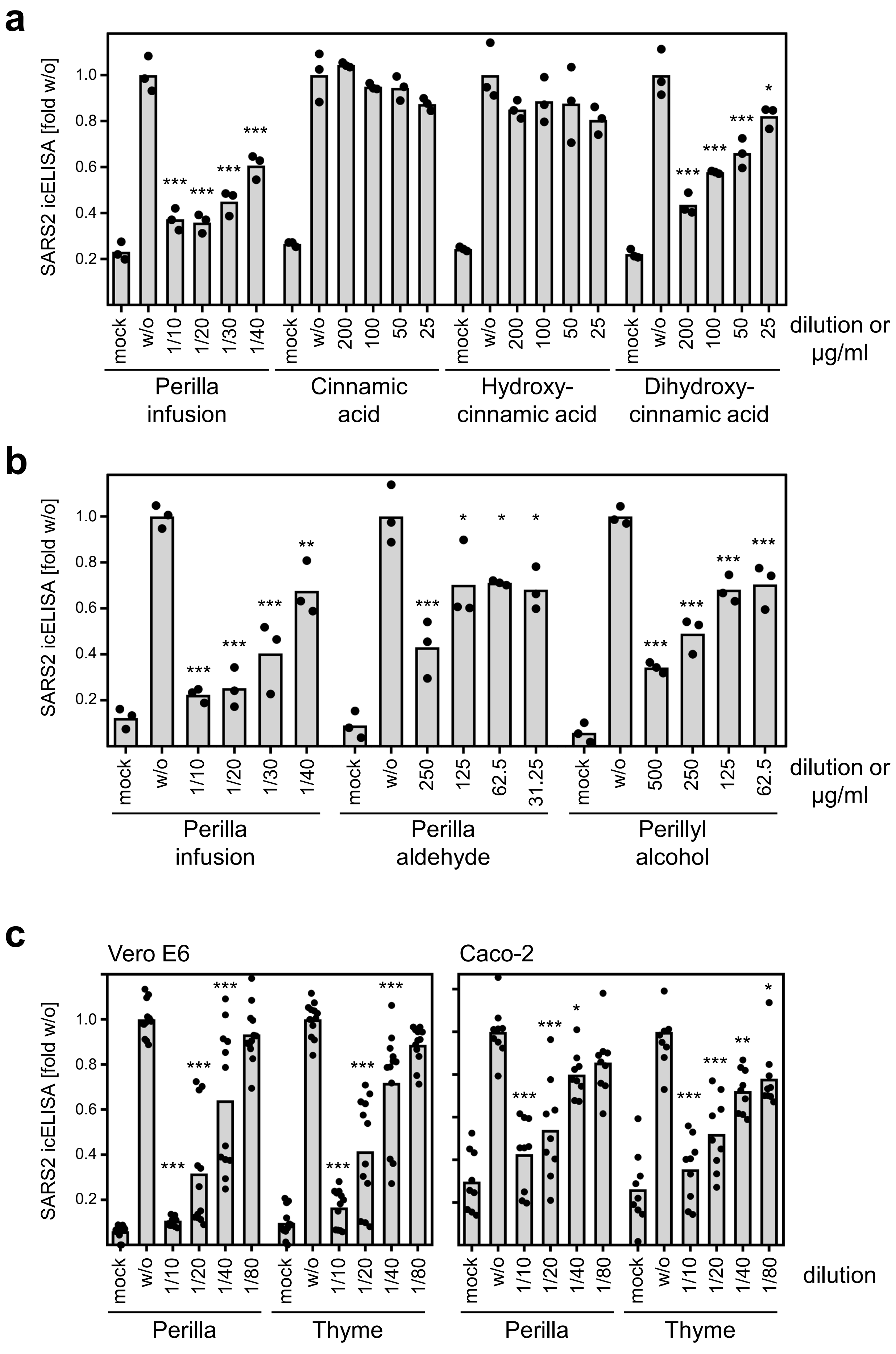

### Supplementary Figure 8

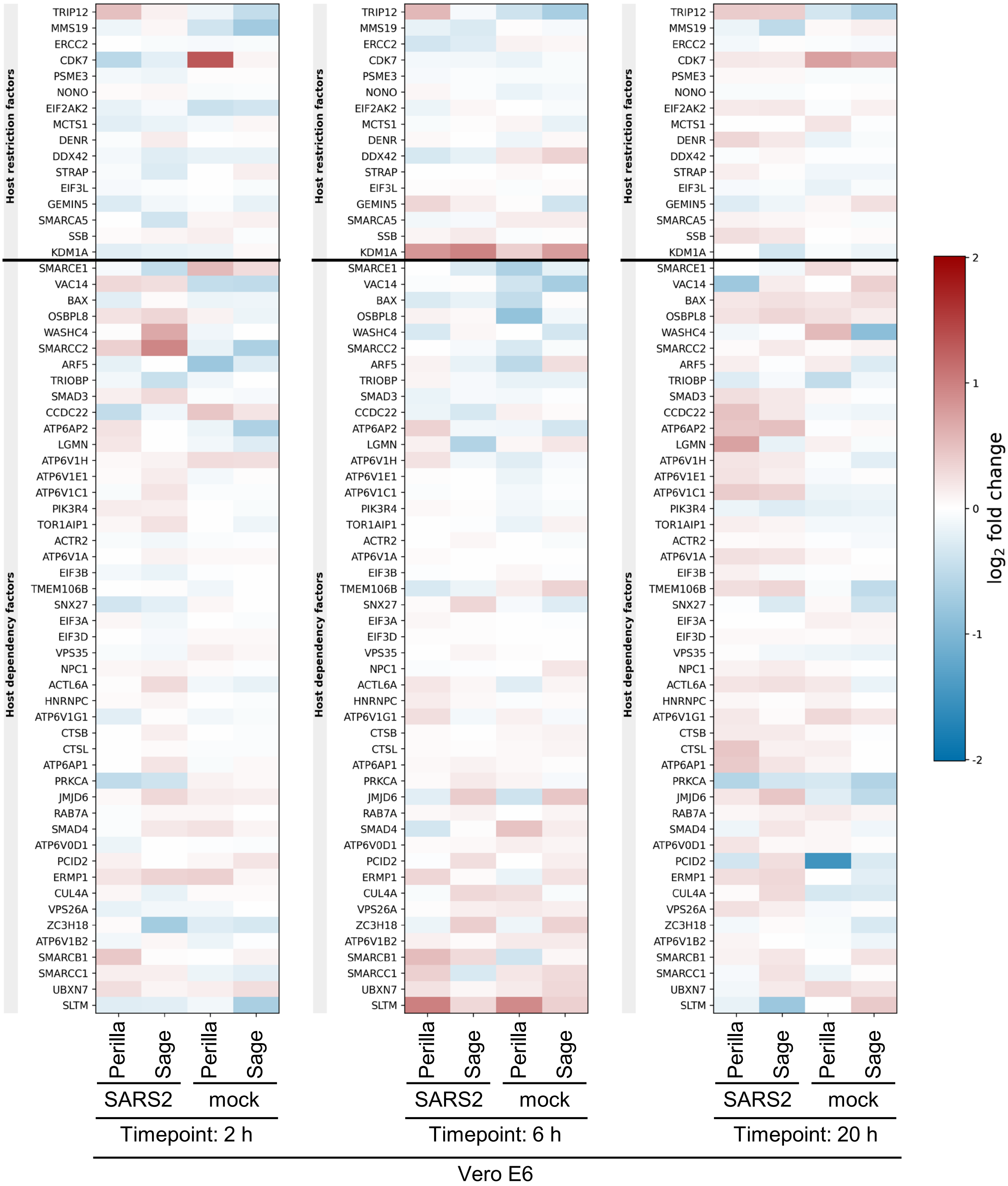

### Supplementary Figure 9

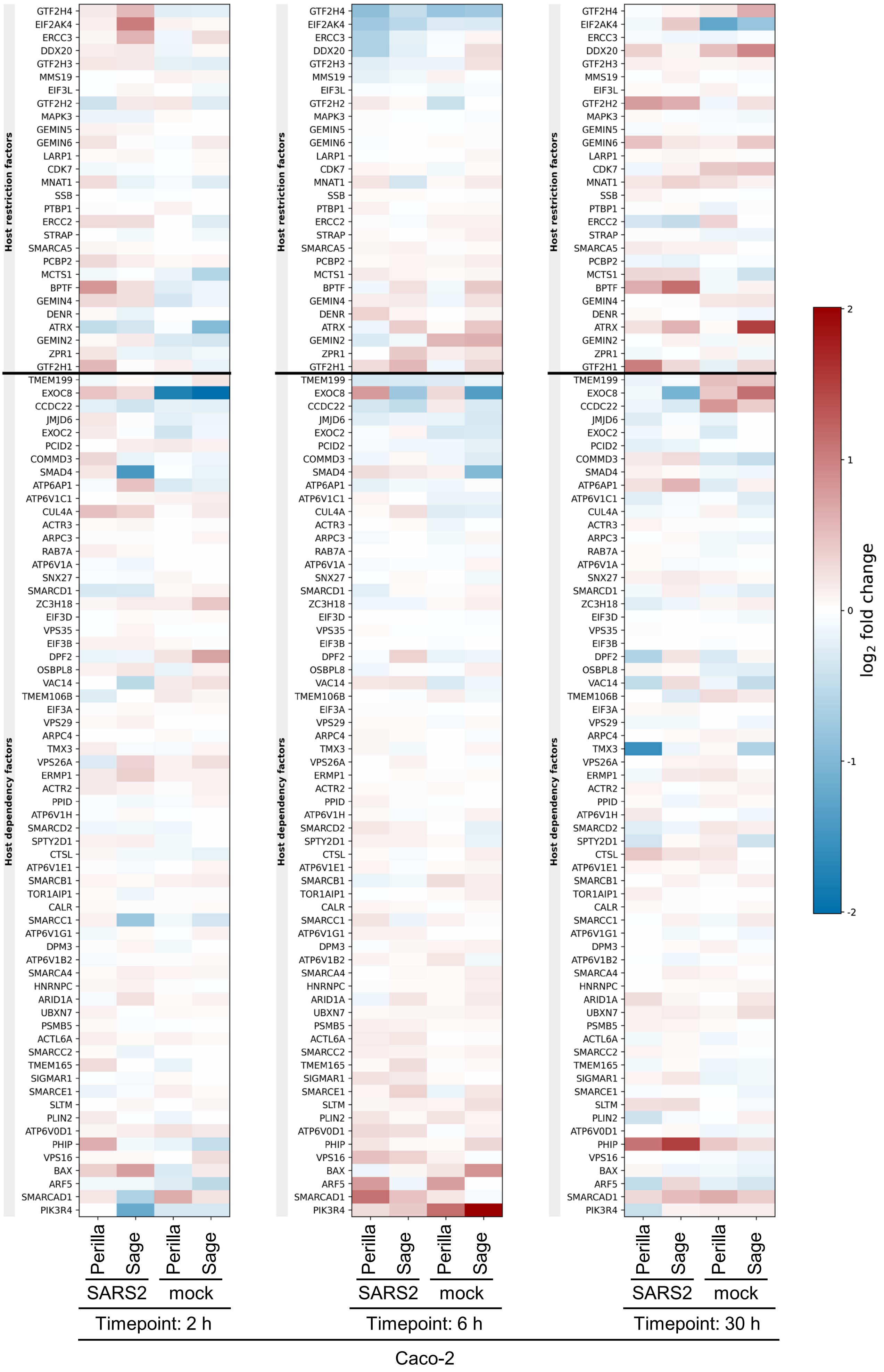

### Supplementary Figure 10

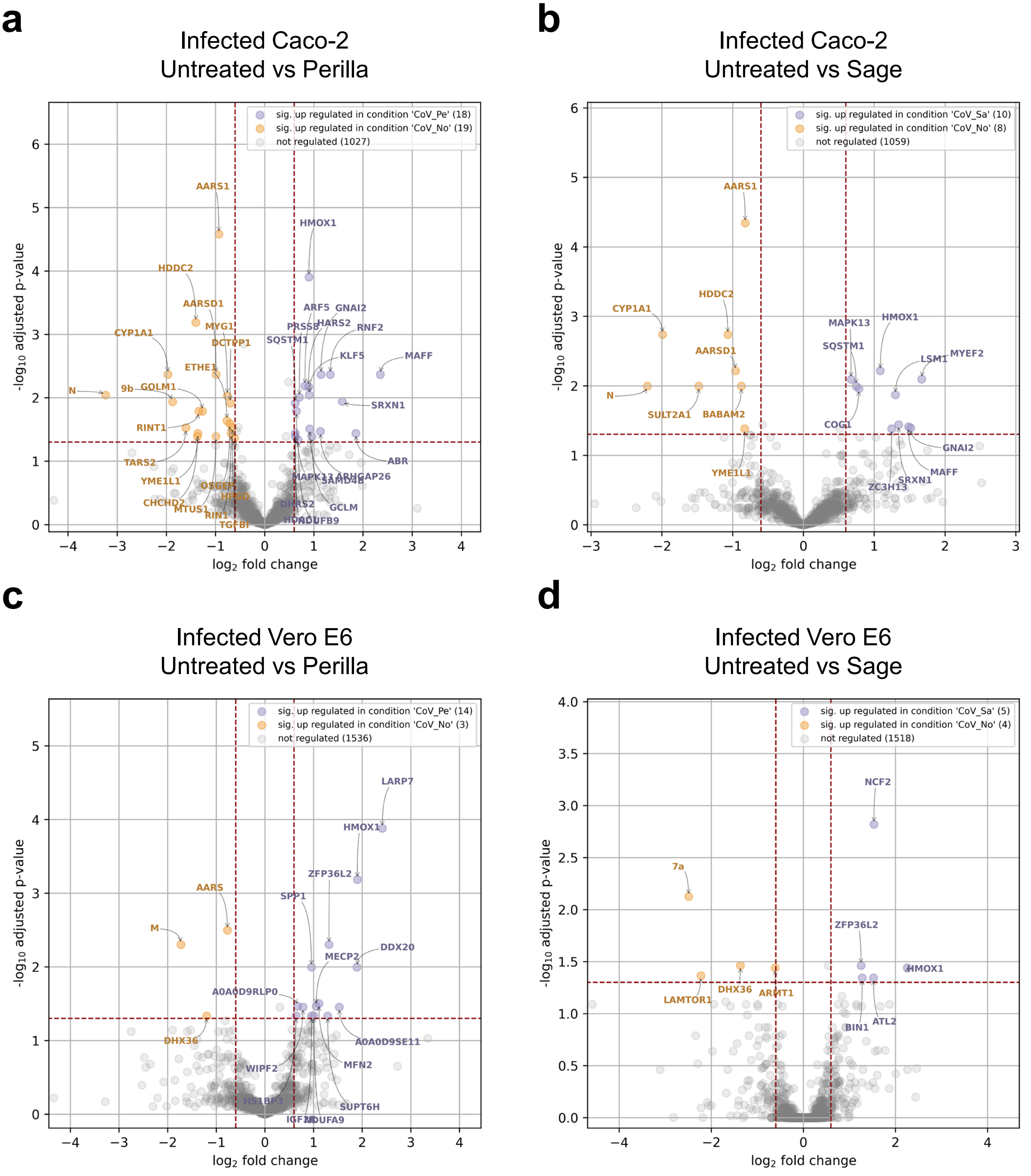

### Supplementary Figure 11

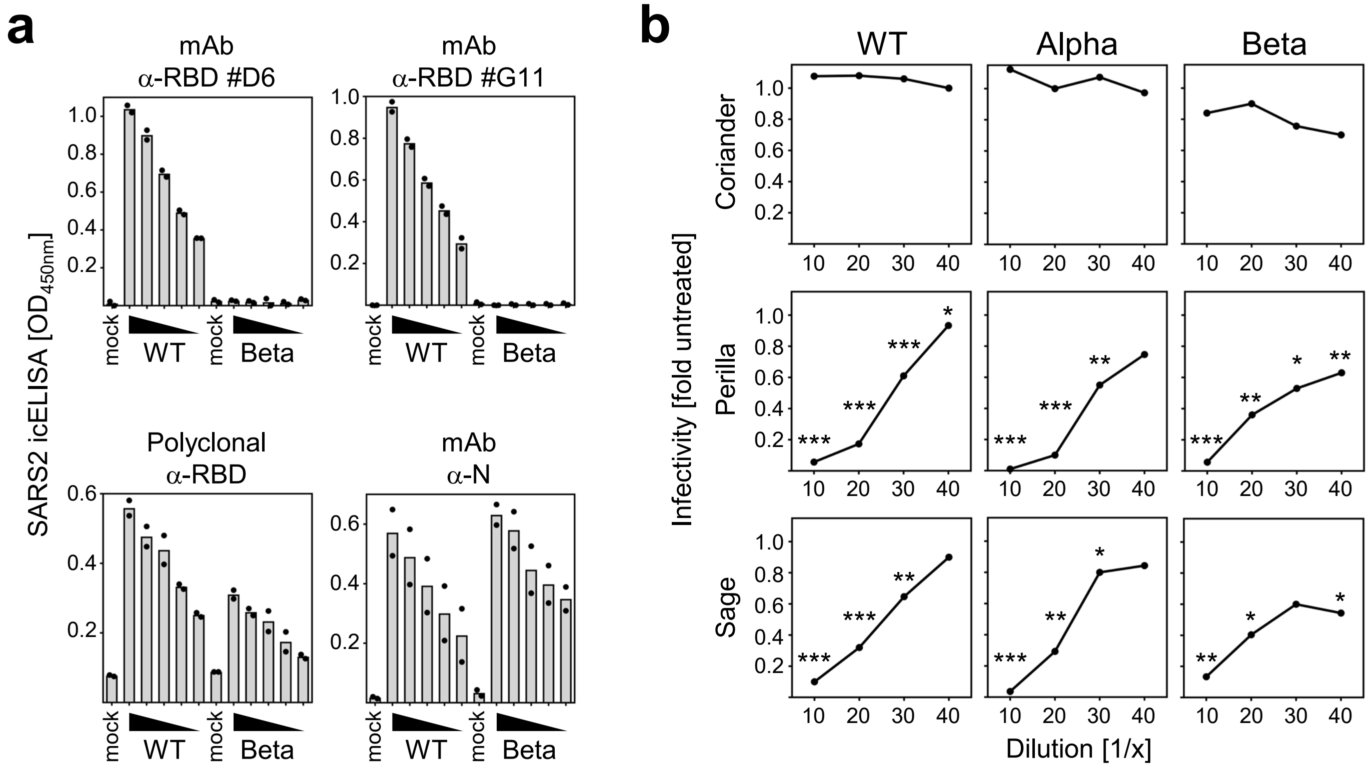
